# Robust estimation of heritability and predictive accuracy in plant breeding: evaluation using simulation and empirical data

**DOI:** 10.1101/671768

**Authors:** Vanda M Lourenço, Joseph O Ogutu, Hans-Peter Piepho

## Abstract

**Background:** Genomic prediction (GP) is used in animal and plant breeding to help identify the best genotypes for selection. One of the most important measures of the effectiveness and reliability of GP in plant breeding is predictive accuracy. An accurate estimate of this measure is thus central to GP. Moreover, regression models are the models of choice for analyzing field trial data in plant breeding. However, models that use the classical likelihood typically perform poorly, often resulting in biased parameter estimates, when their underlying assumptions are violated. This typically happens when data are contaminated with outliers. These biases often translate into inaccurate estimates of heritability and predictive accuracy, compromising the performance of GP. Since phenotypic data are susceptible to contamination, improving the methods for estimating heritability and predictive accuracy can enhance the performance of GP. Robust statistical methods provide an intuitively appealing and a theoretically well justified framework for overcoming some of the drawbacks of classical regression, most notably the departure from the normality assumption. We compare the performance of robust and classical approaches to two recently published methods for estimating heritability and predictive accuracy of GP using simulation of several plausible scenarios of random and block data contamination with outliers and commercial maize and rye breeding datasets.

**Results:** The robust approach generally performed as good as or better than the classical approach in phenotypic data analysis and in estimating the predictive accuracy of heritability and genomic prediction under both the random and block contamination scenarios. Notably, it consistently outperformed the classical approach under the random contamination scenario. Analyses of the empirical maize and rye datasets further reinforce the stability and reliability of the robust approach in the presence of outliers or missing data.

**Conclusions:** The proposed robust approach enhances the predictive accuracy of heritability and genomic prediction while alleviating the need for performing outlier detection for a broad range of simulation scenarios and empirical breeding datasets. Accordingly, plant breeders should seriously consider regularly using the robust alongside the classical approach and increasing the number of replicates to three or more, to further enhance the accuracy of the robust approach.

## Introduction

Genomic studies, whether from an association, prediction or selection perspective, constitute a field of research with increasing statistical methodological challenges given the growing complexity (population structure, coancestry, etc), dimension of datasets, measurement errors and atypical observations (outliers). Outliers often arise from atypical environments, years, field pests or other phenomena. Here, regression models are the tool of choice whether in studies involving human, animal or plant applications. However, it is well known that the performance of these models is poor when their underlying assumptions are violated and their unknown parameters are estimated by the classical likelihood [49]. For example, violation of the normality assumption – depending on its severity – may lead to both biased parameter estimates and coefficients of determination [7] and strongly interfere with variable selection [5]. In the case of the linear mixed model, such violation can tamper with the estimation of variance components [24], which itself can be very challenging even when data are normally distributed but the sample size is small. Violation of model assumptions due to contamination of data with outliers can have several other deleterious effects on regression models. In genomic association studies, for example, departure from normality can induce power loss in the detection of true associations and inflate the number of detected spurious associations [22]. In plant genomics such violations of model assumptions and the associated biases often translate into inaccurate estimates of heritability and predictive accuracy [10]. This can have significant practical consenquences because predictive accuracy is the single most important measure of the performance of genomic prediction (GP). The reduction of these adverse effects through the use of more robust methods is thus of considerable practical importance [48].

Recently, [9] proposed a method for estimating heritability and predictive accuracy simultaneously (Method 5) and compared its performance with several contending methods from the literature including a popular method in animal breeding (Method 7). More details on Methods 5 and 7 can be found in the ‘Genomic Prediction’ Section. The authors concluded from these comparisons that Methods 5 and 7 consistently gave the least biased, most precise and stable estimates of predictive accuracy across all the scenarios they considered. Additionally, Method 5 gave the most accurate estimates of heritability [9]. Both methods are founded on the linear mixed effects model as well as on ridge regression best linear unbiased prediction (RR-BLUP) through a two-stage approach [34–36]. The first stage of this two-stage approach involves phenotypic analysis and thus is likely to be adversely affected by contaminated phenotypic plot data. In particular, contamination can undermine the accuracy with which the adjusted means are estimated in the first stage and thus negatively impact estimation of both heritability (only Method 5) and predictive accuracy in the subsequent second stage where RR-BLUP is used [15]. [10] later examined the performance of the same seven methods in the presence of one outlying observation under 10 simulated contamination scenarios. These simulations reaffirmed that Methods 5 and 7 performed the best overall and produced the best estimates of both heritability (only Method 5) and predictive accuracy across all the contamination scenarios they considered. However, one outlying observation for their dataset with a sample size of 698 genotypes corresponds to a level of contamination of merely 0.1%. As stated by [10], outliers may arise in plant breeding studies from measurement errors, inherent characteristics of the studied genotypes, enviroments or even years. As the process generating the outliers may vary across locations and/or trials, it is conceivable that a non-neglegible percentage of phenotypic observations may be typically contaminated when large field trial datasets are considered. As a result, the composite effects of such substantial levels of contamination on the accuracy of methods for estimating heritability and accuracy of GP can be potentially considerable. Such outliers may not always be easy to detect and eliminate prior to phenotypic data analysis. Therefore, using robust statistical procedures for phenotypic data analysis of field trial datasets can help ameliorate the adverse effects of outliers.

Robust statistical methods have been around for a long time and are designed to be resistant to influential factors such as outlying observations, non-normality and other problems associated with model misspecification [17]. Therefore, the use of robust methods has been advocated for inference in the linear and linear mixed model setups [6, 25], as well as in ridge regression [1, 15, 26, 27, 45, 52]. As a result of such considerations and the recent advances in computing power, it is not surprising that there has been a strong, renewed interest in exploring these techniques to robustify existing methods or develop new procedures robust to moderate deviations from model specifications [24, 41].

Consequently, to tackle the problem of biased estimation of heritability and predictive accuracy due to contamination of phenotypic data with outliers, we aim to robustify the first phase of the two-stage analysis used in GP. Such an approach will, in addition, largely obviate the need to check for and eliminate mild or even extreme outliers from the data prior to analysis. We use a Monte-Carlo simulation study encompassing several contamination scenarios to assess the performance of the proposed robust approach relative to: (i) the approach used by [35], and (ii) simulated underlying true breeding values taken as the gold standard. These assessments are carried out at each of the two stages involved in predicting breeding values by comparing the accuracy with which the two approaches estimate true genotypic values in phenotypic analysis. In a third stage, we compare the heritabilities (H2) and predictive accuracies (PA) estimated by the two competing approaches using Method 5 (H^2^ and PA) and Method 7 (PA only). In addition, we compare the heritability estimated by Method 5 with the generalized heritability estimated by Oakey’s method [29]. The latter method was not evaluated by [9].

Also, an application of the methodology to real commercial maize (*Zea mays*) and rye (*Secale sereale*) datasets is presented and used to empirically assess the usefulness of the proposed robust approach. Lastly, we discuss how to effectively apply the proposed robust approach to phenotypic data analysis and the estimation of heritability and predictive accuracy of GP in plant breeding.

The robust and the classical approaches are implemented in the **R** software using the code in the supplementary materials (Additional file **AppendixE Rcode.pdf**). The ASREML-R package is used to fit the models at the second stage.

## Materials and Methods

### Datasets

#### Rye dataset

The Rye data were obtained from the KWS-LOCHOW project and is described in more detail elsewhere [2, 3]. These data consist of 150 genotypes tested between 2009 and 2011 at several locations in Germany and Poland, using *α* designs with two replicates and four checks (replicated two times in the two replicates). Each trial was randomized independently of the others. The field layout of some trials was not perfectly rectangular. Trials at some locations and for some years had fewer blocks but larger size, i.e., two different sizes were used for a few trials. Blocks were nested within rows in the field layout. The dataset has 16 anomalous observations pertaining to distinct genotypes, that the breeders identified as outliers. Moreover, yield was not observed for one genotype. For this example we consider two complete datasets (320 observations): the first is the original dataset without any corrections, which we call the ‘raw’ dataset, and the second is the original dataset with the 16 yield observations replaced with missing values, which we refer to as the ‘processed’ dataset. In addition, we consider a cleaned version of the raw dataset (288 observations; called cleaned dataset) obtained by removing from the raw data the 16 outlying genotypes (32 observations) identified by both the breeders and the criterion used for outlier detection described in the ‘Example Application’ Section. We note that because the empirical rye dataset has only two replicates, a single outlier will automatically generate an outlier with the same absolute value of opposite sign for the other replicate of the same genotype. Consequently, we removed a testcross genotype entirely from the cleaned dataset even if only one of its two replicate observations was outlying. The raw, processed and cleaned datasets comprise only 148, 148 and 132 genotypes with genomic information, respectively.

#### Maize dataset

The maize dataset was produced by KWS in 2010 for the Synbreed Project. The data set has 1800 yield observations on 900 doubled haploid maize lines and 11,646 SNP markers. Out of the 900 test crosses 698 were genotyped whereas 202 were not. The test crosses were planted in a single location (labelled RET) on nine 10 by 10 lattices each with two replicates. Six hybrid and five line checks connected the lattices (398 observations in total). The lines were crossed with four testers. After performing quality control, the breeder recommended replacement of 38 yield observations with missing values. A more elaborate description of this maize dataset is provided in [9, 11].

For this example we consider two datasets each with 1800 yield observations: the first is the original dataset without any corrections, which we call the ‘raw’ dataset, and the second one is the original dataset with the 38 yield observations replaced with missing values, which we refer to as the ‘processed’ dataset. Furthermore, we consider a third dataset (called cleaned raw dataset) obtained by removing 46 outliers from the raw dataset. The fourth dataset (called the cleaned and processed dataset) is obtained by removing seven outliers from the processed dataset. All the outliers satisfied the criterion for outliers described in the ‘Example application’ Section. As with the rye dataset, we removed a testcross genotype entirely from the raw dataset if at least one of the two replicate observations was outlying. Thus, the raw, processed, cleaned raw and cleaned and processed datasets have 1800, 1754, 1800 and 1793 yield observations and 698, 687, 698 and 697 genotypes with genomic information, respectively.

### Genomic prediction

#### True correlation

The correlation between the true (**g**) and the predicted 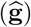 breeding values (true correlation or true predictive accuracy) can be calculated from simulated data as

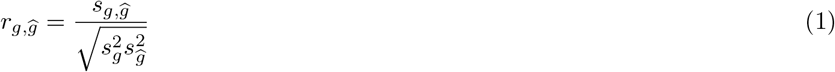

where 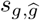 is the sample covariance between the true and predicted breeding values, 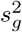 and 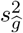 are the sample variances of the true and predicted genetic breeding values, respectively. This correlation is often the quantity of primary interest in breeding studies. The simulation study therefore assesses the accuracy with which 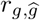 is estimated by Methods 5 and 7, whose details are described below.

#### Two-stage approach for predicting breeding values

[9] use the two-stage approach of [35] to predict true breeding values (**g**) that are then used to estimate heritability and predictive accuracy. This approach is quite appealing because it greatly alleviates the computational burden of the single-stage approach [47], without compromising the accuracy of the results.

The single-stage model can be written as

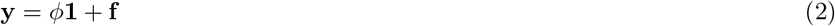

where **y** is the vector of the observed phenotypic plot values, *ϕ* is the general mean, **f** is a vector that combines all the fixed, random design and error effects (*replicates*, *blocks*, etc.). For the simulated data **f** has four random effects only and is given by **f** = **Z**_*g*_**g** + **Z**_*r*_**u**_*r*_ + **Z**_*b*_**u**_*b*_ + **e** where (i) **Z**_*g*_ is the design matrix for the genotypes with 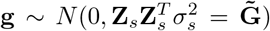, **Z**_*s*_ is the matrix of biallelic markers of the single nucleotide polymorphisms (SNPs), coded as −1 for genotypes `AA`, 1 for `BB` and 0 for `AB` or missing values and 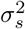 is the variance of the marker effects; (ii) **Z**_*r*_ is the design matrix for the *replicate* effects with 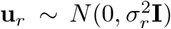 and 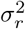 is the variance of the *replicate* effects; (iii) **Z**_*b*_ is the design matrix for the *block* effects with 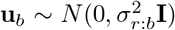 and 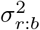 is the variance of the *block* effects; and (iv) **e** ~ *N* (0, **R**) are the residual errors and **R** is the variance-covariance matrix of the residuals. In our model 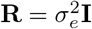 where 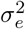 is the residual plot error variance.

The two-stage approach basically breaks this model into two models. In the first stage, which we seek to robustify, we use the model

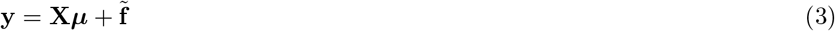

where **y** is defined as before, **X** = **Z**_*g*_ is the design matrix for the genotype means, ***μ*** = *ϕ***1**+**g** is the vector of unknown genotypic means and 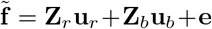. Note that in this first stage the genomic information regarding the SNP markers (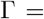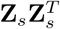) is excluded from this analysis because genotype means ***μ***, which comprise the genetic effects **g**, are modelled as fixed. This is usually the case when stage-wise approaches are considered, in which case the genomic information is included only in the last stage [35].

In the second stage, the genotype means 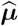 estimated at the first stage are used as a response variable in a model for computing the true breeding values **g** specified as

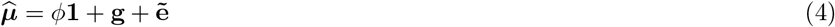

where *ϕ* is the general mean and 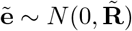 with 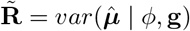.

Note that any standard varieties or checks are dropped from the dataset before the adjusted means 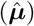 from the first stage are submitted to the second stage. The mixed model equations for (4) can be solved to obtain the best linear unbiased prediction for **g**, 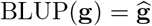, using a ridge-regression formulation of BLUP, i.e., RR-BLUP.

In case weights are used when fitting the second-stage model, then 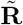 should be replaced by **W**^−1^, with **W** being a weight matrix computed from the estimated first-stage variance-covariance matrix 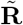. In our case we used Smith’s [46] and standard (ordinary) [35] weights. Specifically, 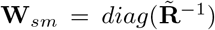 for Smith’s and 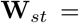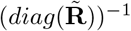 for standard weights, respectively.

More details on the two-stage approach can be found in [9, 35, 36].

#### Method 5

This method (**M**5) calculates predictive accuracy as

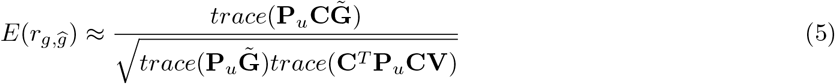

where 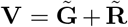 with **V**, 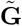 and 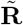 being the variance-covariance matrices for the phenotypes, genotypes and residual errors of the adjusted genotypes, respectively; 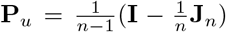, with **J**_*n*_ a *n* × *n* matrix of ones; 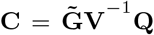, with 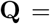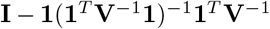, and **1** denoting a vector of ones. Under this formulation, which provides a direct estimate of the correlation between the true (**g**) and the predicted 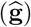 breeding values, the RR-BLUP of **g** is now given by 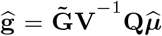 [34].

Heritability can then be computed from (5) as

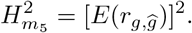

#### Method 7

This method (**M**7) is commonly used by animal breeders to directly compute predictive accuracy (*ρ*) from the mixed model equations (MME, [12, 28, 51]) by firstly computing the squared correlation between the true (**g**) and predicted breeding values 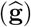, i.e., reliability (*ρ*^2^).

Since the MME for the second-stage model (4) are given by

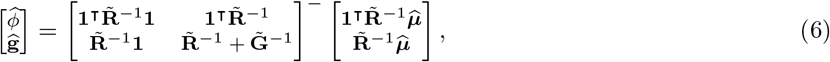

with the variance-covariance matrix of 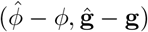 given by

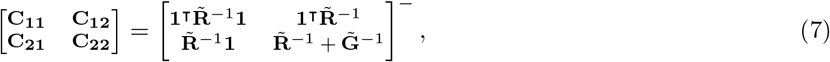

and the variance-covariance matrix of **g** and 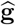 given by

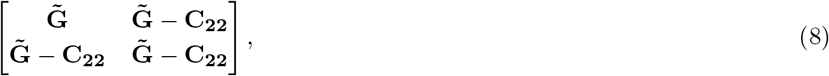

the reliability for each genotype is computed as

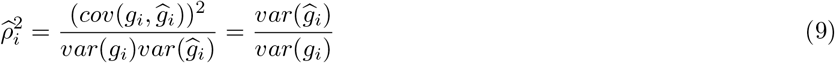

where only the diagonal elements of the matrices 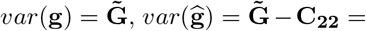 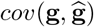 are extracted. The average reliability across the genotypes in each dataset is then estimated by

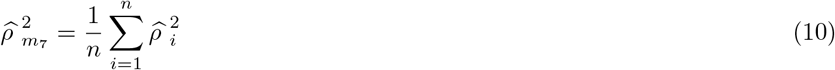

where *n* is the total number of genotypes in the dataset. Predictive accuracy 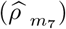 is then computed as the square root of 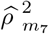. Alternatively, predictive accuracy can be computed as

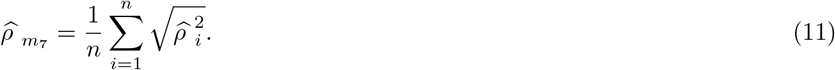

Further details on this derivation can be found in [36].

#### Oakey’s Method

[29] propose a generalized heritability measure that was recently re-expressed by [40] as

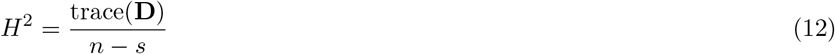

where 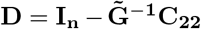 and *s* is the number of zero eigenvalues of **D**. We also use this method to estimate heritability and compare this estimate with the estimate obtained by method **M5**.

### Robust estimation

#### Robust estimation of the linear mixed model for phenotypic data analysis

In this section we briefly review the robust approach of [19] to linear mixed effects models that we use in an attempt to robustify the first stage of the two-stage approach to genomic prediction in plant breeding. This approach is implemented in the R software package *robustlmm* via the function `rlmer()` [20, 21].

We consider the general linear mixed model

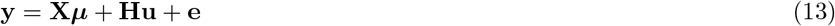

where **y** is a vector of observations, **X** is the design matrix for the fixed effects (intercept included), ***μ*** is the vector of unknown fixed effects, **H** is the design matrix for the random effects, **u** ~ *N* (**0**, **U**) is the vector of unknown random effects and **e** ~ *N* (**0**, **R**) is the vector of random plot errors. Note that for our first-stage model **Hu** = **Z**_*r*_**u**_*r*_ + **Z**_*b*_**u**_*b*_ and ***μ*** = *ϕ***1** + **g**.

Model (13) also assumes that *cov*(**u**, **e**) = 0 and as such we have that

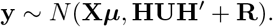

We henceforward assume for simplicity that 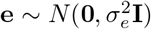 and 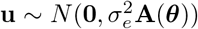 where the variance matrix **A** of the random effects depends on the vector of unknown variance parameters ***θ*** (this assumption can be relaxed to obtain more general formulations, see e.g., [19]). The variance of **y** now simplifies to

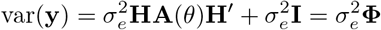

with **Φ** = **HA**(*θ*)**H**′ + **I**.

Because **A**(***θ***) is a positive-definite symmetric matrix and assuming that ***θ*** is known, one can obtain its Cholesky decomposition as chol(**A**(***θ***)) = **B**(***θ***), set **u** = **B**(***θ***)**b** and rewrite model (13) as

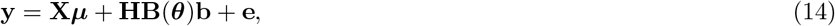

where 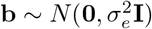 so that we again have 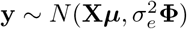.

The classical log-likelihood for (14) can be written as

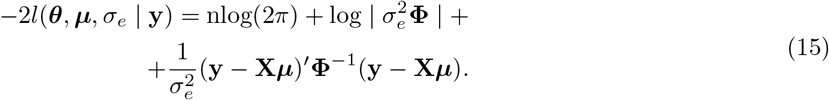

Furthermore, for a given set of ***θ***, ***μ*** and *σ*_*e*_ (44, Chapter 7)

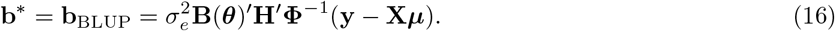

From (15) and (16), an objective function that incorporates the observation-level residuals and the random effects as separate additive terms can be derived and expressed as

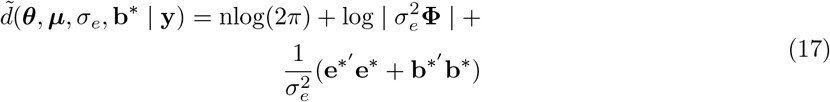

where

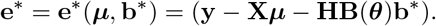

This particular trick is crucial in order to independently control contamination at the levels of the residual and random effects.

Assuming ***θ*** and *σ*_*e*_ are known and taking the partial derivatives of (17) with respect to ***μ*** and **b**^*^, we get the following estimating equations for these effects,

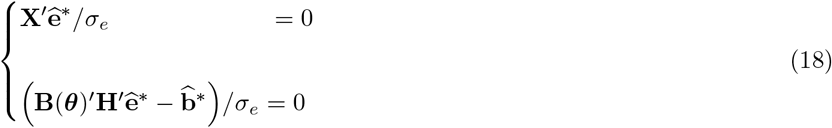

where

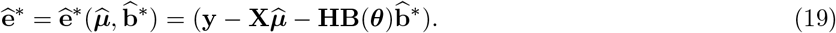

If **B**(***θ***) is diagonal, as in our case, these equations are robustified by replacing 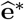 and 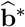 by bounded functions 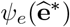 and 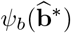, where the *ψ*_*e*_ and *ψ*_*b*_ functions need not be the same:

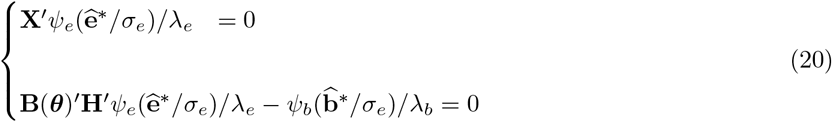

where 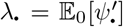 is required to balance the 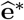 and 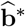 terms in case different *ψ* functions are used; 1/λ_*e*_ and 1/λ_*b*_ are scaling factors (as in M-regression [17]) and cancel out in the special case where *ψ*_*e*_ ≡ *ψ*_*b*_.

If we let

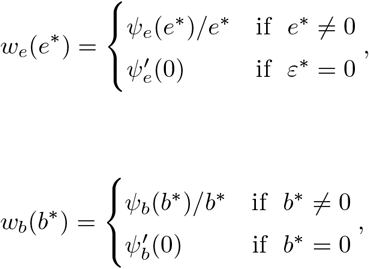

Λ_*b*_ = λ_*e*_/λ_*b*_, 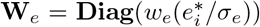 and 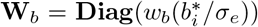, and after some simplification, equation (20) can be written as

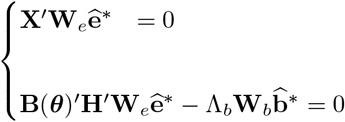

which, after expanding 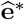 with (19), yields the following system of linear equations:

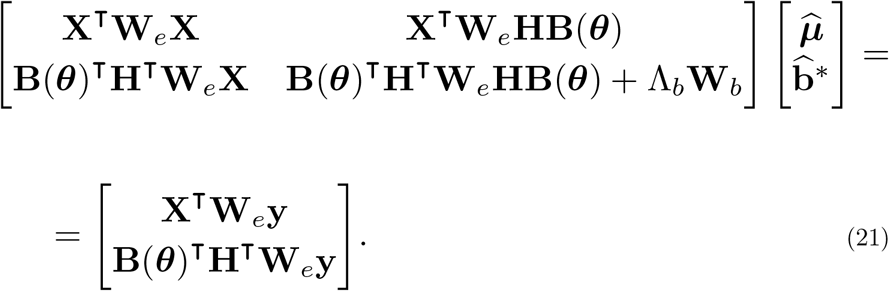

The algorithm for estimating parameters of (21) begins with a predefined set of weights. It then alternates between computing 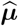 and 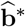 for a given set of weights and updating the weights for a given set of estimates. [18] and [19] provide more details on the estimation of the scale and covariance parameters and the estimation procedure for the non-diagonal case.

If *replicate* and *block* (nested within *replicates*) are the only random effects apart from the residual error in the first-stage model (this is the case for the simulation study for our first-stage model and for the first-stage model for the rye dataset) then 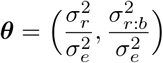, where 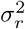 and 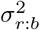 are the variances for the *replicate* and *block* random effects, respectively. Also here, **A**(***θ***) is a two-block diagonal matrix (*k* = 2 blocks). Furthermore, because we assume 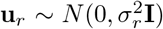 and 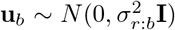 for the first-stage model, **B**(***θ***) = [**A**(***θ***)]^1/2^ is a diagonal matrix.

In particular, for the simulated data consisting of 698 observations of maize *yield* from 2 *replicates* each having 39 *blocks* (more details in the ‘Simulation’ Section), we compute 2 + 39 = 41 weights (**W**_*b*_) for the observations at the level of the random effects and 2 × 698 = 1396 weights (**W**_*e*_) for the observations at the level of the fixed effects (i.e., for the residuals).

#### Robust approach to phenotypic analysis

Phenotypic data derived from field trials are prone to several types of contamination that may range from measurement errors, inherent characteristics of the genotypes and the environments to the years in which the trials were conducted. As such, if contaminated observations are present in the vector of phenotypes **y** in the first stage of phenotypic data analysis, then they can unduly influence the estimation of the means for the testcross genotypes (***μ***) in model (3), resulting in inaccurate estimates of adjusted phenotypic means 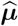. In turn, these possibly inaccurate estimates of ***μ*** are passed on to the second stage of the procedure (model (4); adjusted RR-BLUP) from which the breeding values **g** are estimated. The possibly biased estimates of (**g**) may undermine the accuracy of the estimated heritability and predictive accuracy.

To minimize bias in the estimation of heritability and predictive accuracy, we propose using the preceding robust model for the first stage of phenotypic data analysis. The second stage then proceeds in the same way as the classical method except that, now, the robust estimates 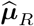 from the first stage are used in (4).

## Simulation

### Simulated datasets

We consider a real maize dataset from the Synbreed Project (2009 − 2014). This dataset was extracted for one location from a larger dataset and consists of 900 doubled haploid maize lines, of which only 698 testcrosses were genotyped, and 11, 646 SNP markers. Six hybrid checks and five line checks were considered and genotypes were crossed with four testers as explained in more detail in [9]. Variance components estimated from this dataset 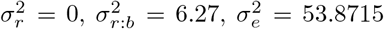 and 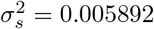 were used to simulate the block and plot effects based on an *α*-design (31) with two replicates and the model

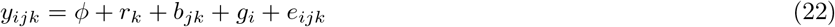

where *y*_*ijk*_ is the yield of the *i*-th genotype in the *j*-th block nested within the *k*-th complete replicate, *ϕ* is the general mean, *r*_*k*_ is the fixed effect of the *k*-th complete replicate, *b*_*jk*_ is the random effect of the *j*-th block nested within the *k*-th complete replicate, *g*_*i*_ is the random effect of the *i*-th genotype, and *e*_*ijk*_ is the residual plot error associated with *y*_*ijk*_. More details on (22) can be found in Table S3 in the supplementary materials of [10].

Our simulations consider 1000 simulated Maize datasets described as follows: each dataset consists of 698 observations of yield in 2 replicates, with the 698 genotypes distributed over 39 blocks as in Table 1. Four out of the 39 blocks have 17 observations, whereas the remaining 35 have 18 observations.

**Table 1:** A sample simulated Maize dataset

| $I$ | rep | block | genotype | yield |
| --- | --- | --- | --- | --- |
| 1 | 1 | 1 | 267 | 7.416505 |
| 2 | 1 | 1 | 149 | 1.945098 |
| . | . | . | . | . |
| . | . | . | . | . |
| . | . | . | . | . |
| 698 | 1 | 39 | 459 | 25.097810 |
| 699 | 2 | 1 | 604 | 12.640605 |
| . | . | . | . | . |
| . | . | . | . | . |
| . | . | . | . | . |
| 1396 | 2 | 39 | 614 | 18.859413 |

### Simulation of outliers

In order to simulate outliers, a percentage of phenotypic observations in the dataset is chosen and contaminated by replacing the observed value of each selected observation by that value plus 5−, 8− or 10− times the standard deviation of the residual error (*σ*) used to simulate the phenotypic datasets. Additionally, we also consider two distinct types of data contamination:

i. Random contamination: 1, 3, 5, 7 and 10% of the phenotypic data in only one of the two replicates are randomly contaminated, amounting to an overall data contamination rate of 0.5, 1.5, 2.5, 3.5 and 5%, respectively.
ii. Block contamination: phenotypic data in 1, 2, 3, 4 and 5 whole blocks in only one of the two replicates are contaminated, amounting approximately to 1.3, 2.6, 3.9, 5.2 and 6.5% overall rate of data contamination, respectively.

We use the notation “%_sd” to denote the random contamination scenarios corresponding to the contamination of a particular percentage (%) of the data with outliers of size `sd` and “block_sd” to refer to block contamination scenarios corresponding to the contamination of a specific number of whole blocks (`block`) with outliers of size `sd`.

### First- and second-stage models

In the first stage (eq.3), we consider *yield* as the response variable, the *genotypes* as the fixed effects and the *replicates* and *blocks nested within replicates* as the random effects. In the second stage (eq.4), we consider the adjusted genotypic means estimated in the first stage as the response variable, the *intercept* as the fixed effect and the *genotypes* as the random effects with a variance-covariance structure given by the genomic relationship matrix.

### Comparing performance of the classical and robust approaches

The performance of the classical and robust approaches is evaluated in three steps, labelled L1, L2 and L3. L1 involves a comparison of results from the first stage; L2 entails a comparison of results from the second stage and L3 focuses on a comparison of the estimated heritability and predictive accuracy, which can be viewed as constituting the third stage. For each of the three levels, we consider the null scenario (uncontaminated datasets), random and block contamination scenarios.

Additionally, the influence of the Smith’s and standard weighting schemes used in the second stage of the two-stage approach are considered in L2.

The following quantities are computed and used to compare the performance of the classical and robust approaches at levels L1–L3.

#### L1

The mean squared deviation (MSD) of the estimated from the true genotypic means is computed for both the classical and robust approaches as

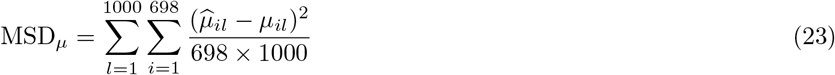

where *μ*_*il*_ is the true mean of the *i*-th genotype in the *l*-th simulation run and 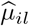 is its estimate.

The estimates of 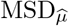 for the classical (*C*) and robust (*R*) approaches are compared for each scenario using

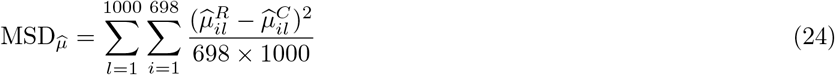

and are expected *a priori* to agree for the null scenario.

It is also instructive to compute and plot

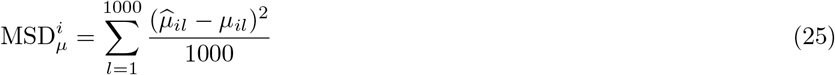

for each genotype *i* = 1, …, 698 for both approaches. Furthermore, the overall estimated genotypic mean (across genotypes and simulations) is also computed and compared to the corresponding true genotypic mean. Moreover, since the rank order of genotypes is also of great importance in plant breeding studies, the Pearson correlation coefficient (*r*_*p*_) between the true and estimated genotypic means (predictive accuracy) is also computed and compared between the two approaches. This yields an estimate of the predictive accuracy for the genomic means.

#### L2

At this level, we compute the MSDs for the genomic breeding values **g** analogously to equations (23)–(25). The *r_p_* between the true and estimated breeding values is again computed and used to compare the two methods and assess any improvement in the estimation of **g** when genomic information is included in the analysis. This provides an estimate of the accuracy of genomic prediction.

#### L3

Here, the methods are compared by computing the following MSDs,

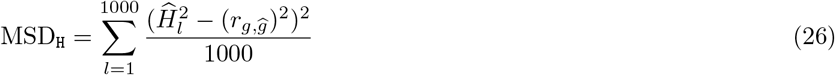

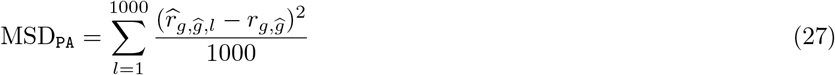

where 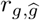 is the Pearson correlation computed between the true and the estimated breeding values and averaged across the 1000 simulations, 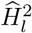 and 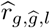 are, respectively, the heritability and predictive accuracy estimated in the s-th simulation via the methods described earlier. These MSDs quantify the deviation of the estimated from the true heritability (H^2^) or predictive accuracy (PA). In addition, we provide boxplots of the estimated heritablity and predictive accuracy for the 1000 simulation runs for each scenario.

## Simulation results

### Null scenario

#### L1

The following computed MSDs

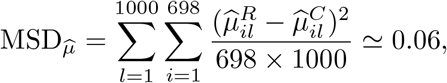

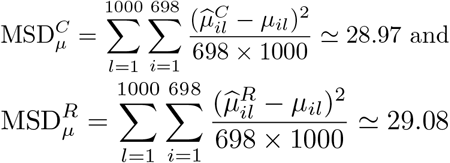

show, as expected, that both methods perform similarly when the data are not contaminated 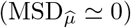. However, the classical method performs slightly better than the robust one 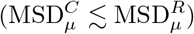. Even so, both MSD values are not particularly close to zero. Still, as these MSD values are squared deviations averaged across all the 1000 simulation runs and 698 genotypes, they seem reasonable.

The slightly better performance of the classical relative to the robust method is also apparent in the per-genotype MSDs (Figure 1S). The two approaches produce virtually identical estimates for the overall mean of 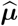 (i.e., mean{*μ_il_*}, *i* = 1, …, 698, *l* = 1, …, 1000) and *r_p_* (Table 2).

**Table 2:** Estimated overall mean of 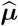 and predictive accuracy expressed as the Pearson correlation coefficient *r_p_* obtained using the classical (CLS) and the robust (ROB) methods (averaged across the 1000 simulations)

| true mean=8.923 | ROB | CLS |
| --- | --- | --- |
| overall mean $\hat{\mu}$ | 8.906 | 8.908 |
| $r_p$ | 0.764 | 0.765 |

**Figure 1:**
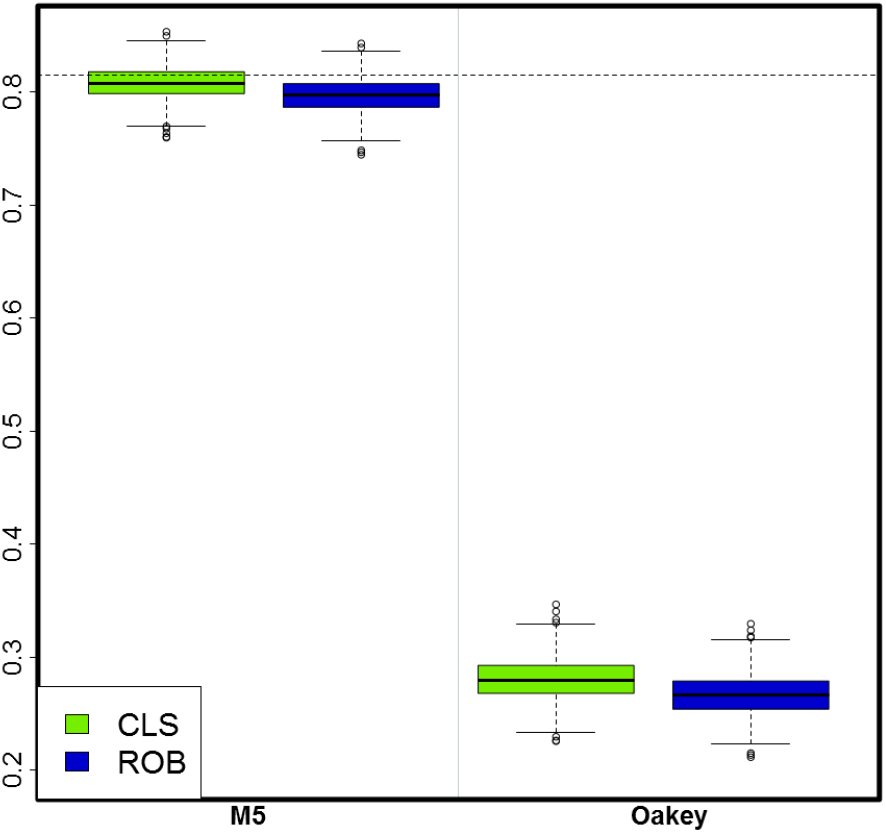
Estimates of heritability obtained from methods **M5** and Oakey’s for the classical (CLS) and robust (ROB) approaches

The two methods estimate the variances of both the random effects and the residual errors equally well (Figure 2S).

**Figure 2:**
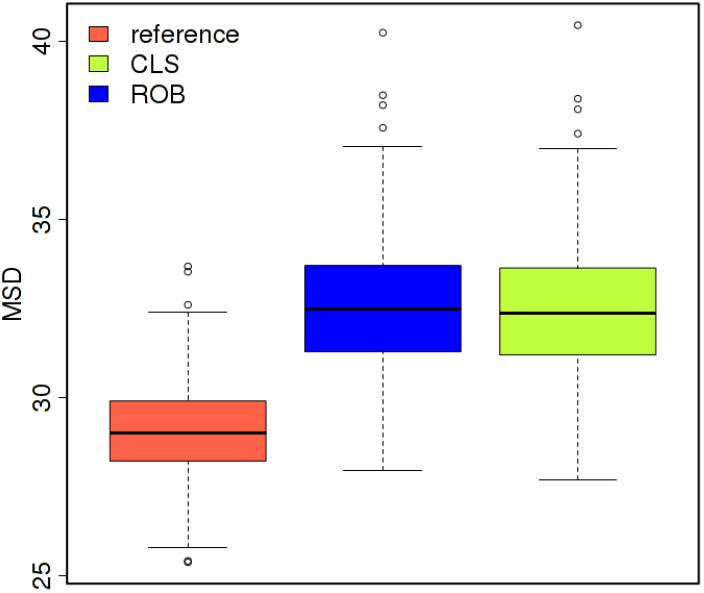
Boxplot of the 698 per-genotype mean squared deviations 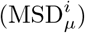 under the null scenario (classical as reference) against the ones obtained in the random scenarios 1_5 for the classical (CLS) and robust (ROB) approaches

The Smith’s and standard weights obtained in the first stage for both the classical and robust approaches are very small (Figure 3S). Precisely, the MSD between the two different types of weights is approximately 0.6 × 10^−6^ and the MSD between the values of each type of weight computed by the two approaches is about 0.6 × 10^−5^.

**Figure 3:**
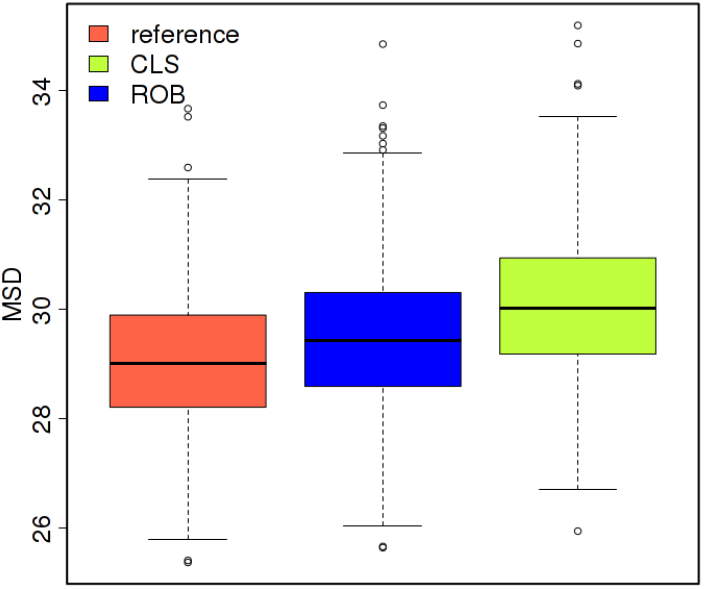
Boxplot of the 698 per-genotype mean squared deviations 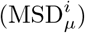 under the null scenario (classical as reference) against the ones obtained in the block scenarios 1_5 for the classical (CLS) and robust (ROB) approaches

#### L2

There were no major differences between the estimated breeding values obtained using either the standard or Smith’s weighting schemes at the second stage (MSD_*g*_ ≃ 25 for both cases). For this reason we only present results produced using Smith’s weights.

The MSDs for the second stage

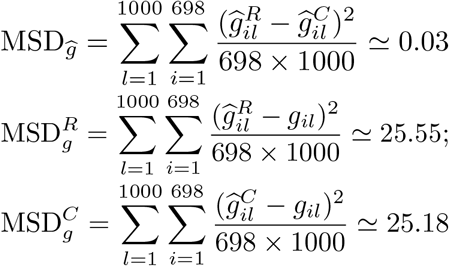

show a modest improvement over the corresponding estimated genotypic means at the first stage and that the methods continue to perform similarly as in the first stage. Relative to the estimates for the first stage, the per-genotype MSDs (Figure 13S) increase for about 22% but decreases for about 47% of the genotypes. This trend is similar for both the classical and robust approaches. Additionally, for the second stage, the mean *r_p_* = 0.903 for both approaches. This increase in *r_p_* relative to the first stage (≃ 18.2%) shows that using genomic information at the second stage improves genomic prediction and hence the ranking of genotypes. For the overall mean of the EBVs 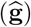, it drops to ≃5 from ≃9 for both approaches (first row, Tables 2S & 4S).

Quite interestingly, in terms of the estimation of the genetic variance, the robust approach performs slightly better than the classical (Figure 14S).

#### L3

Both the classical and robust approaches produce the following MSDs for heritability (Method **M**5 only) and predictive accuracy (Methods **M**5 and **M**7):

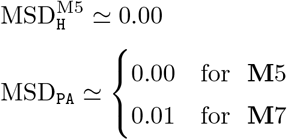

showing the estimates of heritability and predictive accuracy to be quite accurate. We note that estimates of heritability and predictive accuracy were computed by fixing the residual variance from the first stage to one as described in the ‘Genomic prediction’ Section. In general this produced more accurate estimates than the alternative for which the residual variance estimated in the first stage is used. Therefore all the results displayed here for the third stage use the former implementation.

Boxplots for the estimated PA (methods **M5** and **M7**) and H^2^ (method **M5** only) across the 1000 simulations for the null scenario are shown together with the ones for the random and block contamination scenarios (Figures 19S-20S). These suggest that method **M**5 produces more accurate estimates of PA than method **M**7.

Relative to method **M**5, Oakey’s heritability estimates are unacceptably lower than the simulated true values (Figure 1). To further explore why these estimates were remarkably smaller than those produced by **M**5 or the simulated true values, we took 100 out of the 1000 simulation replicates and refitted the entire two-stage model by setting 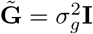 at the second stage. The heritability estimates for M5 and Oakey’s were virtually identical (MSD≃ 0.1 × 10^−27^). This strongly suggests that Oakey’s method works fine with independent genotypes but performs poorly when the model used to estimate heritability has a kinship matrix. Consequently, we do not consider Oakey’s heritability estimates further except in a few comparisons in the ‘Example application’ Section.

### Random contamination scenarios

#### L1

The MSD_*μ*_ for these scenarios are similar between approaches for each level of contamination and size of outlier (Tables 3 and 1S). Hence for the random contamination scenarios, the robust and classical approaches produce comparable estimates for the genotypic means. The per-genotype MSDs also reaffirm the similar performance of the two approaches (Figure 4S). Nevertheless, it is noteworthy that even for the least extreme scenario, 1% contamination with an outlier of size 5 sd, the Figure 1: Estimates of heritability obtained from methods **M5**and Oakey’s for the classical (CLS) and robust (ROB) approaches increase in the per-genotype 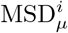 is non-negligible relative to the corresponding values computed by the classical method under the null scenario and used as a benchmark (Figure 2). The per-genotype MSDs increase greatly with increase in the percentage contamination and size of outliers (Figure 4S).

**Table 3:** MSD_***μ***_ between the estimated genotypic means and the true breeding values considering the classical (CLS) and robust (ROB) approaches and the random contamination scenarios.

| Random scenarios |  | CLS | ROB |
| --- | --- | --- | --- |
| % cont | sdt |  |  |
| 0 | - | 28.97 | 29.08 |
| 1 | 5 | 32.44 | 32.51 |
| 1 | 8 | 37.83 | 37.78 |
| 1 | 10 | 42.80 | 42.64 |
| 5 | 5 | 46.26 | 46.39 |
| 5 | 8 | 72.98 | 72.87 |
| 5 | 10 | 97.50 | 97.25 |
| 10 | 5 | 63.30 | 63.65 |
| 10 | 8 | 116.40 | 116.67 |
| 10 | 10 | 165.22 | 165.30 |

The estimated overall mean of the estimated genotypic means 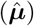 for both approaches departs increasingly from the true overall mean of ***μ*** as both the level of contamination and size of outliers increase (Table 2S). The same trend is evident for the *r_p_*, implying deterioration of the ranking of genotypes (Table 2S).

The two methods also differ with respect to how well they estimate particular variance components. More precisely, the classical method estimates the variance for *blocks nested within replicates* 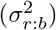 somewhat better than the robust method does from 5% contamination upwards. However, the robust method estimates the variances for *replicates* 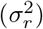 and *residual errors* 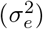 far better than the classical method does (Figures 6S–8S).

The Standard and Smith’s weights computed for both the classical and robust approaches across the random contamination scenarios are shown in Figures 9S and 10S.

As the percentage of contamination and size of the outliers increase, the degree of overlap of the empirical frequency distributions of the classical and robust weights evidently reduces. In particular, the distributions do not overlap at all from the 3% contamination level upwards for the 8− and 10−sd shift-outliers. Also, the weights show an overall decreasing trend, which is more evident for the classical approach and the Standard weights.

#### L2

The mean squared deviations (MSD_**g**_) between the EBVs and the TBVs for the classical approach are displayed in Table 5S. The MSD_**g**_’s for the Standard and Smith’s weights show some minor differences in favour of the latter from 7% contamination and 8-sd shift-outliers onwards. Thus, only the second-stage results obtained using the Smith’s weights are presented in the remainder of this section.

The robust approach tends to produce smaller MSDs between the EBVs and the TBVs as the percentage of random contamination and the size of the shift-outliers increase (Tables 4 & 1S). The second-stage per-genotype MSDs do not show the increasing trend observed for the per-genotype MSDs from the first stage with values ranging between 0 and 100 (Figure 16S). In addition, the robust method always produces higher estimated predictive accuracy, expressed as the averaged Pearson correlation coefficient *r_p_*, than the classical method, implying better ranking of the genotypes (Table 2S). The overall mean EBVs 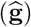 is similar for both approaches but drops steadily as the percentage contamination and size of outliers increase, implying underestimation.

**Table 4:** Mean squared deviation of the estimated from the true genomic breeding values (MSD_**g**_) for the classical (CLS) and robust (ROB) approaches under the random contamination scenarios.

| Random scenarios |  | CLS | ROB |
| --- | --- | --- | --- |
| % cont | sdt |  |  |
| 0 | - | 25.18 | 25.55 |
| 1 | 5 | 26.29 | 26.26 |
| 1 | 8 | 27.84 | 26.44 |
| 1 | 10 | 29.16 | 26.49 |
| 5 | 5 | 29.37 | 28.72 |
| 5 | 8 | 34.53 | 29.97 |
| 5 | 10 | 38.31 | 30.33 |
| 10 | 5 | 32.53 | 32.41 |
| 10 | 8 | 40.39 | 37.40 |
| 10 | 10 | 45.71 | 39.04 |

The robust approach also produces more accurate estimates of the marker-effect variance 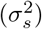 up to 10% contamination and 5-sd shift-outliers (Figure 15S). For the 10% contamination scenarios with 8- and 10-sd shift-outliers the robust estimates of 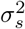 no longer overlap with the true marker-effect variance but their boxplots show a smaller inter-quartile range (IQR) and lower dispersion than those for the classical approach.

#### L3

The robust approach produced generally more accurate estimates for both H2 and PA for the random contamination scenarios (Figures 18S-20S). However, both approaches tend to underestimate both parameters as the percentage contamination and size of outliers increase. The 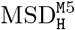 ranged between approximately 0.00 − 0.09 and 0.00 − 0.07 for the classical and robust methods, respectively. The corresponding, 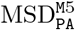 ranged between approximately 0.00 − 0.02 and 0.00 − 0.01 whereas 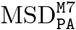 ranged between approximately 0.01 − 0.04 and 0.01 − 0.07. Overall, method **M5**performs somewhat better than method **M7**in estimating predictive accuracy (Figures 18S-20S; Table 6S).

### Block contamination scenarios

#### L1

Although the MSD_*μ*_ for the block contamination scenarios is relatively stable for the robust approach (between 29.08 and 30.11), it increases with increasing level of contamination and size of the outliers for the classical approach (Tables 5 & 3S). In the worst block contamination scenario (5 10) the MSD for the classical approach is about 1.7 times larger than that for the robust approach. The per-genotype MSDs show even poorer performance for the classical method in estimating each of the 698 genotypic means (Figure 5S). By contrast, the robust approach maintains the errors at roughly the same level across the contamination scenarios; a level that is close to the one estimated for the null scenario. This is an attractive property of this method.

**Table 5:** Mean squared deviation of the estimated genotypic means from the true breeding values (MSD_***μ***_) for the classical (CLS) and robust (ROB) approaches under the block contamination scenarios.

| Block scenarios |  | CLS | ROB |
| --- | --- | --- | --- |
| No. blocks | sdt |  |  |
| 0 | - | 28.97 | 29.08 |
| 1 | 5 | 30.90 | 29.44 |
| 1 | 8 | 30.75 | 29.69 |
| 1 | 10 | 31.16 | 29.82 |
| 3 | 5 | 32.06 | 29.64 |
| 3 | 8 | 35.22 | 29.96 |
| 3 | 10 | 38.02 | 30.11 |
| 5 | 5 | 35.31 | 29.76 |
| 5 | 8 | 43.33 | 29.83 |
| 5 | 10 | 50.66 | 29.88 |

**Table 6:** Mean squared deviation of the estimated from the true breeding values (MSD_**g**_) for the classical (CLS) and robust (ROB) approaches under the block contamination scenarios.

| Block scenarios |  | CLS | ROB |
| --- | --- | --- | --- |
| No. blocks | sd |  |  |
| 0 | - | 25.18 | 25.55 |
| 1 | 5 | 25.35 | 25.61 |
| 1 | 8 | 25.38 | 25.63 |
| 1 | 10 | 25.39 | 25.63 |
| 3 | 5 | 25.34 | 25.65 |
| 3 | 8 | 25.37 | 25.68 |
| 3 | 10 | 25.37 | 25.67 |
| 5 | 5 | 25.39 | 25.58 |
| 5 | 8 | 25.40 | 25.58 |
| 5 | 10 | 25.40 | 25.59 |

Block contamination had generally less debilitating effect on the accuracy of the estimated genotypic means than random contamination (Tables 1S & 3S). For example, the block contamination scenario 1_5, which corresponds to an overall 1.3% data contamination, produces smaller MSDs than the random contamination scenario 1_5, which corresponds to only 0.5% overall contamination (Figures 2 and 3; Table 5)

The performance of the two methods also differed noticeably with respect to the estimation of the overall mean of 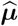. For example, for the worst case scenario (block 5 10) the overall mean of 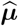 deviated from the true mean by merely 5.5% for the robust approach but by 50.2% for the classical approach (Table 4S), indicating superior performance of the robust approach. Nevertheless, the poor predictive performance of the classical approach at the first stage does not necessarily translate to a reduced predictive accuracy *r_p_* because it does not alter the relative ranking of the genotypes (Table 4S). Accordingly, the ranking of the genotypes does not differ much between the two approaches (estimated *r_p_* ≃ 0.76 for both approaches across all scenarios).

An overall superior performance of the robust compared to the classical approach is also evident for the accuracy of the estimated variance components (Figures 6S-8S).

#### L2

In this case, the MSD_**g**_ obtained in the second stage differ depending on whether the Smith’s or the standard weights are used. In particular, using the Smith’s weights produces more stable MSD_**g**_ estimates across all the *block* contamination scenarios than using the standard weights, which tend to increase with increasing number of contaminated *blocks* and size of outliers (Table 5S). For this reason, only results obtained using the Smith’s weights are presented in the remainder of this section.

For all levels of contamination and size of outliers, the robust overall MSDs between the EBVs and the TBVs did not differ much and fluctuated around ≃ 25 (Table 5S), a value that is similar to the corresponding value for the null scenario (Table 6).

The per-genotype MSD_*g*_ values vary little with increasing size of outliers but suggest that the classical method performs slightly better than the robust method (Figure 17S).

The average estimated predictive accuracy (*r_p_*) across all scenarios was approximately 0.90 for both approaches (Table 4S). Predictive accuracy thus increased from the first to the second stage for the classical (by 17%) and robust (by 18%) approaches, an increase comparable to that observed under the null scenario.

Finally, the robust method estimates the marker-effect variance 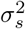 more accurately than the classical method throughout all the block contamination scenarios (Table 15S).

#### L3

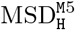 and 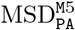 were both ≃ 0.00 for both the classical and robust approaches across all the block contamination scenarios, with the classical producing marginally better results than the robust approach (Figures 18S and 19S). 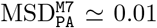 for both approaches with the robust estimates of PA obtained via **M**7 showing slightly greater dispersion (Figure 20S). It is noteworthy that estimates of H^2^ and PA are rather stable across block contamination scenarios (Figures 18S-20S), consistent with the estimated marker-effect variances (Figure 15S).

## Example application

In this Section, we comparatively evaluate differences in the performances of the classical and robust approaches on raw empirical rye and maize datasets prior to quality control. Substantial differences in results between the two approaches would imply problems with the data that require closer inspection by the breeder or data analyst. Such inspection can be followed by data cleaning, which can be a very challenging and time-consuming task. For the two example datasets in this section, we perform data cleaning based on a simple rule of thumb that relies on the weight given to each observation by the robust method. Specifically, observations assigned weights smaller than 0.5 are flagged as outliers. More sophisticated outlier detection techniques are outside the scope of this paper [3, 23]. We apply the classical and robust approaches to the cleaned dataset and compare the results with each other and with the results for the raw dataset. We note that cleaning the data does not necessarily make it conform to model assumptions such as the normality of the errors. We note further that because empirical datasets for both examples each have only two replicates, the robust method usually assigns the same, or very similar, weights to both replicates. This is the reason that a testcross genotype is removed entirely from the cleaned dataset even if only one of its two replicate observations is outlying. This problem would be eliminated by replicating each genotype three or more times.

We similarly analyze the empirical datasets after taking into account the recommendations of the breeder based on quality control to demonstrate that quality control alone will not always detect and eliminate all sources of data contamination and hence does not preclude the use of robust statistical methods.

### Rye dataset

In this example we consider only one trial from the Rye dataset described in the ‘Materials and methods’ Section, which otherwise has the same structure as the simulated maize data set shown in Table 1. The first- and second-stage models fitted to the Rye data set are the same as those described in the ‘Simulation’ Section.

The classical and robust approaches produced strikingly different estimates for the residual and blocks variances at the first stage as well as for heritability and predictive accuracy at the third stage (Table 7; CLS^*r*^ and ROB^*r*^ results). The robust weights assigned to each of the 320 observations in the first stage identified 32 observations for the exact same 16 genotypes identified as outliers by the breeders. When the 32 observations are removed from the data, which amounts to around a 10% reduction in the size of the dataset, then the classical and robust approaches produce very similar estimates, as is expected when data conform to the model’s assumptions (Table 7; CLS^*r*^* and ROB^*r*^* results). In particular, the distributions of the residuals from the classical first- and second-stage models fit to the cleaned dataset satisfy the normality assumption (Shapiro-Wilk normality test: p= 0.9771 and p= 0.6974, respectively), but the distribution of the residuals from the raw dataset do not (Shapiro-Wilk normality test: p< 10^−9^). Inspection of the QQ plots of the residuals (not shown) further reinforced the results of the normality tests. This example clearly demonstrates how the robust approach ameliorates most of the devastating influences of outliers on the classical method. Thus, contamination with outliers inflates the estimated residual variance ~ 20 times for the classical method but only ~ 3.6 times for the robust method. By contrast, contamination reduces the estimated block variance from ~ 11.5 to zero for the classical method but from ~ 11.9 to ~ 8.2 for the robust method. Lastly, contamination reduces the estimated heritability and predictive accuracy far more strikingly for the classical than for the robust approach. However, contamination inflates the marker-effect variance equally by a factor of two for both approaches. Although far lower than the simulated true values, Oakey’s heritability estimates are also shown and compared between the full and the cleaned datasets for completeness (Table 7).

**Table 7:** The estimated *residual* 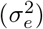, *replicate* 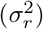 and *blocks within replicates* 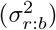 variance components, *genetic* variance 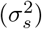, heritability computed by Method **M5** (H2.M5) and predictive accuracy computed by Methods **M5** (PA.M5) and **M7** (PA.M7), using the classical (CLS) and robust (ROB) approaches, for the rye dataset.

| Stage | Parm<sup>†</sup> | CLS<sup>r</sup> | ROB<sup>r</sup> | CLS<sup>qc</sup> | ROB<sup>qc</sup> | CLS<sup>r*</sup> | ROB<sup>r*</sup> |
| --- | --- | --- | --- | --- | --- | --- | --- |
| **1** | $\sigma_e^2$ | 120.1400 | 17.6797 | 5.5267 | 5.3369 | 5.5267 | 5.3674 |
| | $\sigma_r^2$ | 4.4262 | 6.4327 | 5.2124 | 5.7228 | 5.2124 | 5.7276 |
| | $\sigma_b^2$ | 0.0000 | 8.1911 | 11.5441 | 11.8846 | 11.5441 | 11.8804 |
| **2** | $\sigma_s^2$ | 41.2101 | 40.4765 | 24.6136 | 24.3552 | 21.8182 | 21.5183 |
| **3** | H2.M5 | 0.4628 | 0.7387 | 0.8373 | 0.8339 | 0.8305 | 0.8259 |
|  | H2.OK | 0.1906 | 0.4681 | 0.6220 | 0.6160 | 0.6173 | 0.6094 |
|  | PA.M5 | 0.6803 | 0.8594 | 0.9150 | 0.9132 | 0.9113 | 0.9088 |
|  | PA.M7 | 0.6531 | 0.8428 | 0.9038 | 0.9018 | 0.8997 | 0.8969 |

<sup>†</sup> Parm=Parameter;
<sup>r</sup> refers to the raw dataset, i.e, the original dataset before quality control;
<sup>qc</sup> refers to dataset after quality control; <sup>r\*</sup> refers to the cleaned raw dataset;

Results for the processed dataset are similar for the two approaches (Table 7; CLS^*qc*^ and ROB^*qc*^). They are also quite similar to the results for the cleaned dataset, except for the estimated marker-effect variances, which are smaller for the cleaned dataset perhaps due to the decrease in the sample size at the second stage. Interestingly, for this particular case, the residuals from the first stage of the classical model fit satisfy the normality assumption (Shapiro-Wilk normality test: p= 0.6974) but those from the second stage only marginally pass the normality test (Shapiro-Wilk normality test: p= 0.0437). A quick look at the QQ plot of these residuals reveals two residuals that deviate substantially from the equality line (plot not shown). The robust method did not, however, assign any observation a weight smaller than 0.5 and and hence we did not analyse the cleaned and processed datasets. Nevertheless, if a less conservative threshold of 0.7, say, were used instead of 0.5, then, the robust method would have flagged one observation for a check genotype and two for one testcross genotype.

### Maize dataset

In the first stage (eq.3), we consider *yield* as the response variable, the *genotypes* as the fixed effects and the *trials*, the *replicates nested within trials* and the *blocks nested within replicates nested within trials* as the random effects. In the second stage (eq.4), we consider the adjusted genotypic means estimated in the first stage as the response variable, the *intercept* as the fixed effect and the *genotypes* as the random effects with a variance-covariance structure given by the genomic relationship matrix. Note that only the 698 genotypes with available genomic information are submitted to the second stage. In addition, 46 observations of yield (amounting to around 2.6% overall contamination) were identified as outliers by the robust weights computed from the robust first-stage model using the raw dataset. Here, all the observations assigned weights of 0.5 or less by the robust model were classified as outliers. Among these 46 outliers, 22 corresponded to 11 genotypes with genomic information, meaning that the second stage for the cleaned raw dataset comprised only 687 (698 − 11) genotypes. Of the remaining 24 outliers, 18 correspond to 9 genotypes with no genomic information and 6 to 3 hybrid checks and 3 line checks. Overall, 11 + 9 = 20 test crosses and 4 of the 6 checks are a subset of the 38 yield observations recommended for removal (deletion) by the breeder during quality control. The robust method identified only 7 observations as outliers from the processed maize dataset (i.e., with 38 missing yield observations). Two of the 7 observations came from one genotyped test cross, 2 hybrid and 3 line checks. Furthermore, two out of these 7 outliers were also identified when the raw dataset was analysed with the robust method. A detailed treatment of outlier detection strategies is beyond the scope of this paper and can be found elsewhere [4, 23, 39, 48].

As with the Rye dataset, the classical and robust approaches produced different results for the full dataset (Table 8; CLS^*r*^ and ROB^*r*^ results). The differences between the two methods at the first and second stages translate into major differences in the estimated heritability and predictive accuracy. For the cleaned dataset (Table 8; CLS^*r*^* and ROB^*r*^* results) the methods produce more similar estimates of variance components, heritability and predictive accuracy, although estimates are not as close as the ones observed in the case of the rye dataset. In addition, the robust results for the full dataset are close to those obtained via the classical method applied to the cleaned dataset. Note that removal of the outliers was sufficient for the residuals from the first stage but not the second stage of the classical model fit to conform to the normality assumption, in contrast to the results for the rye dataset.

**Table 8:** The estimated *residual* 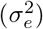, *trial* 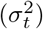, *replicates within trials* 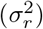 and *blocks within replicates within trials* 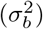 variance components, *genetic* variance 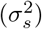, heritability computed by Method **M5** (H2.M5) and predictive accuracy computed by Methods **M5** (PA.M5) and **M7** (PA.M7), using the classical (CLS) robust (ROB) approaches, for the maize dataset.

| Stage | Parm<sup>†</sup> | CLS<sup>r</sup> | ROB<sup>r</sup> | CLS<sup>qc</sup> | ROB<sup>qc</sup> | CLS<sup>r*</sup> | ROB<sup>r*</sup> | CLS<sup>qc*</sup> | ROB<sup>qc*</sup> |
| --- | --- | --- | --- | --- | --- | --- | --- | --- | --- |
| 1 | $\sigma_e^2$ | 352.4962 | 56.0533 | 47.8927 | 49.1810 | 46.5067 | 48.6339 | 45.0147 | 47.9068 |
| | $\sigma_t^2$ | 7.5289 | 9.7426 | 6.3445 | 10.5256 | 9.1892 | 12.0945 | 8.7991 | 12.2285 |
| | $\sigma_r^2$ | 3.4126 | 3.7156 | 3.4067 | 2.9704 | 3.2630 | 3.1115 | 3.6797 | 3.0902 |
| | $\sigma_b^2$ | 0.0000 | 4.8448 | 7.9319 | 6.6856 | 8.0854 | 6.5774 | 7.7984 | 6.4579 |
| 2 | $\sigma_s^2$ | 0.0008 | 0.0067 | 0.0074 | 0.0058 | 0.0096 | 0.0071 | 0.0085 | 0.0062 |
| 3 | H2.M5 | 0.2573 | 0.8020 | 0.8328 | 0.8013 | 0.8634 | 0.8231 | 0.8504 | 0.8099 |
|  | H2.OK | 0.0219 | 0.2732 | 0.3139 | 0.2713 | 0.3671 | 0.3035 | 0.3420 | 0.2824 |
|  | PA.M5 | 0.5073 | 0.8960 | 0.9126 | 0.8951 | 0.9292 | 0.9077 | 0.9222 | 0.8999 |
|  | PA.M7 | 0.4705 | 0.8184 | 0.8338 | 0.8178 | 0.8486 | 0.8287 | 0.8426 | 0.8221 |

<sup>†</sup> Parm=Parameter; <sup>r</sup> refers to the raw dataset, i.e, the original dataset before quality control;
<sup>qc</sup> refers to dataset after quality control; <sup>r\*</sup> refers to the cleaned raw dataset;
<sup>qc\*</sup> refers to the cleaned 'after quality control' dataset.

The results for the full dataset after quality control (Table 8; CLS^*qc*^ and ROB^*qc*^) are similar to those from the robust method (ROB^*r*^; Table 8) However, the residuals from the classical first-stage model fit violate the normality assumption. After removing the 7 outliers from the processed dataset (CLS^*qc*^* and ROB^*qc*^*) the classical and robust approaches produced even more similar results (Table 8) but the residuals still depart from the normality assumption.

As before, Oakey’s heritability estimates are also provided and compared between the full and the cleaned datasets (Table 8).

## Discussion

The simulation results showed that the classical and robust approaches perform similarly when datasets are not contaminated and thus conform to the linear model assumptions. This is a desirable property for any method that seeks to be an alternative to the classical method. Since datasets do not usually conform to all model assumptions, we assessed the relative performance of both methods in estimating genetic breeding values, heritability and predictive accuracy, across a range of contamination scenarios with outliers, which tamper mostly with the assumptions of normality and variance homogeneity of the residuals. All the scenarios involved either random or block contamination (mimicking plausible field conditions), and for each contamination type, differed only in the percentage of the observations that were contaminated and the size of the outliers. Also, two weighting schemes were used with each dataset in the second stage of the two-stage approach.

The simulations revealed that block contamination has a lesser impact on the estimated parameters than random contamination. Also, the estimated true breeding values improve from the first to the second stage, based on the Pearson correlation coefficient, reaffirming the value of using genomic information in the analyses. In addition, the use of the Smith’s weights produces more consistent parameter estimates from the second stage onwards and is therefore recommended for the two-stage approach.

A comparison of the performance of the classical and robust two-stage approaches is summarized in Table 7S. In general, the proposed robust method shows a superior performance to the classical approach. In terms of the accuracy of heritability and genomic prediction, the robust approach clearly outperforms the classical for the random contamination scenarios but performs similar performance to the classical approach for the block contamination scenarios. Also, method **M**5 produces more accurate estimates of predictive accuracy of genomic prediction than method **M**7. Quite surprisingly, the simulations suggest that Oakey’s method is unsuitable for estimating heritability when using a model with a kinship matrix.

Interestingly for the block contamination scenarios, the robust method generally outperformed the classical in both the first and second stages, but this did not translate into higher predictive accuracies. This is likely because the block effect (i.e., effect of blocks within replicates) is completely confounded with the effect of contamination within blocks. As a result, if the block effect is included in the model at the first stage it captures the effect of contamination within the block, yielding an inflated block variance for the classical but not the robust approach. This explains why the performance of the classical approach improves from the first to the second stage. It also emphasizes the need to include a random *block* effect in the first stage to account for intrablock variance especially when using the classical approach.

A noteworthy observation from the simulations is that if a study design has only two replicates, then the robust or the classical methods cannot identify only one of the two replicate observations as an outlier. Hence, using an automated cleaning process, one has to discard twice as many observations as the actual number of outliers. This is because given only two replicates, a single outlier results in two large residuals with the same absolute value but opposite signs. This makes it hard to determine which of the two replicates is actually the outlier.

The robust method can also be useful to breeders doing variety testing for which only the first-stage model is required. Here, the robust approach had clearly superior performance for the block contamination scenarios. For the random contamination scenarios, except for the *blocks within replicates* variance, the robust method produced more accurate estimates of the variance components than the classical method did. Moreover, because late-generation breeding trials typically use only two replicates as breeders aim to maximize the number of different environments, the robust method will merely downweight but not require deleting both replicate observations if it identifies either one or both of them as outliers. This property of the robust method is highly desirable because it enables the plant breeder to obtain genomic predictions for all the target genotypes. By contrast, using the classical method only would result in discarding all the genotypes for which at least one of the two replicate observations is an outlier. This is because it is impossible to determine which of the two replicate observations is the true outlier.

The analysis of the real datasets also furnished some insights about the performance of the methods. For the rye dataset, for example, the 16 outliers identified by the breeders, were also detected by the robust model. Each of the 16 outliers belonged to a distinct block, thus mimicking a random contamination scenario. Also, 8 outliers were observed in the first replicate and the remaining 8 on the second replicate hence resulting in a balanced distribution of outliers at the replicate level. The differences between the results obtained by the classical and robust approaches for the complete dataset are consistent with the ones observed for the simulated random contamination scenarios. Removing the outliers from the data produced a closer agreement between the classical and the robust results but some slight difference from the results produced by the robust approach for the complete dataset still remained. A plausible explanation for this is that removing outliers from the data may: (i) substantially reduce sample size; (ii) alter the distribution of the data and (iii) potentially lead to the underestimation of variances for the cleaned data. This last point precisely matches what is observed for the estimated residuals and marker-effect variances in the two empirical data analyses.

The first-stage results from the analysis of the full empirical raw maize dataset showed a huge discrepancy in the estimated *residual* variance component, moderate disagreement in the estimated *block* variances and similar estimates of *replicate* and *trial* variances between the classical and robust approaches. This result is surprising and deviates from expectation based on the results of the analyses of the simulated data sets. A possible explanation for this unexpected result may relate to the difference in the models fitted to the simulated data and the empirical maize data set and the nature of the outliers. In this case, the 46 observations removed were unevenly spread across 17 out of the 20 blocks (of size 90) and amount to 3% of all the data. Two of these 17 blocks had approximately 8 − 9% contamination. The criterion used to identify the outliers in the maize data set was the robust weights computed from the robust model fit and was somewhat conservative as only the observations assigned weights equal to 0.5 or less were flagged. This criterion, when applied to the rye dataset, correctly identified the 16 outliers that had already been identified by the breeders. However, for the maize dataset this approach to outlier identification is probably too restrictive because the distribution of the residuals from the classical first-stage model fit to the cleaned dataset satisfied the normality assumption but the residuals from the classical second-stage model fit did not. This observation reinforces the view that successfully cleaning the data to eliminate outliers prior to analysis, plus satisfactorily addressing the drawbacks listed above can be exceedingly challenging. Of the 38 yield observations replaced with missing values as recommended by the breeder based on quality control, 24 were identified by the robust method as outliers based on the analysis of the raw dataset and consisted of either negative or zero yield values, which are evidently anomalous. The other 14 of the 38 deleted observations were plausible and were not identified as outliers by the robust method. Results of the analysis of the processed maize dataset with 38 missing yield observations set to missing were very similar between the two approaches. In particular, the results are also quite similar to those from the analysis of the raw dataset using the robust method. This finding emphasizes the stability and reliability of the robust approach both in the presence of outliers and missing observations.

## Conclusion

In conclusion, we show not only the advantages of a robust approach to phenotypic data analysis and genomic prediction but also provide new insights into the potential problems associated with using the classical approach to phenotypic data analysis and genotypic prediction in plant breeding. The proposed robust approach, enhances the accuracy of genomic prediction while alleviating the need for performing outlier detection. Accordingly, plant breeders would do well to seriously consider using these robust methods regularly alongside the classical approach.

## Acknowledgements

This work received partial financial support from Portuguese National Funds by the Fundação para a Ciência e a Tecnologia (Portuguese Foundation for Science and Technology) through Projects PTDC/MAT-STA/0568/2012 and UID/MAT/00297/2013 (CMA: Centro de Matemática e Aplicações) and Sabbatical Grants SFRH/BSAB/105935/2014 and SFRH/BSAB/142919/2018. VML and HPP acknowledge partial support by the Conselho de Reitores das Universidades Portuguesas (CRUP) and the Deutscher Akademischer Austauschdienst (DAAD) through the Luso-German Integrated Actions Project PT/A13/17-DE/57339863. HPP acknowledges support by the German Research Foundation (DFG, Grant PI 377/18-1). JOO was supported by the German Research Foundation (DFG, Grant OG 83/1-1).

We thank KWS LOCHOW GMBH for providing the rye dataset and KWS for providing the maize dataset. We also thank Paul Schmidt for the helpful discussion regarding Oakey’s heritability results.

## Availability of data and materials

Additional file SupplMaterial.pdf refers to: (i) supplementary tables and figures that are referenced in the manuscript; (ii) additional simulation results; (iii) an R code to simulate a synthetic dataset, which we call toydata, that includes the simulation of phenotypic data as well as SNP marker data and kinship; (iv) R code to run the robust two-stage approach.

