## Supplementary material for "Robust estimation of heritability and predictive accuracy in plant breeding: evaluation using simulation and empirical data": Suppl Tables and Figures

### Appendix A

#### Supplementary tables

Table 1S. Mean squared deviations of the estimated genotypic means from the true breeding values – $\text{MSD}_\mu = \sum_{i=1}^{698}\sum_{j=1}^{1000}\frac{(\widehat{\mu}_{ij}-\mu_{ij})^2}{698\times 1000}$ – and mean squared deviations of the estimated breeding values from the true breeding values – $\text{MSD}_g = \sum_{i=1}^{698}\sum_{j=1}^{1000}\frac{(\widehat{g}_{ij}-g_{ij})^2}{698\times 1000}$ – for the classical (CLS) and robust (ROB) methods under the **random** contamination scenarios. MSDs of the classical from the robust estimates are also reported (CLS-ROB).

| **Scenarios I** | | **1st-stage ($\widehat{\boldsymbol{\mu}}$)** | | | **2nd-stage ($\widehat{\mathbf{g}}$)** | | |
|---|---|---|---|---|---|---|---|
| % cont | sdt | **CLS** | **ROB** | **CLS-ROB** | **CLS** | **ROB** | **CLS-ROB** |
| 0 | - | 28.97 | 29.08 | 0.06 | 25.18 | 25.48 | 0.05 |
| 1 | 5 | 32.44 | 32.51 | 0.09 | 26.29 | 26.26 | 0.13 |
| 1 | 8 | 37.83 | 37.78 | 0.15 | 27.84 | 26.44 | 0.54 |
| 1 | 10 | 42.80 | 42.64 | 0.21 | 29.16 | 26.49 | 1.01 |
| 3 | 5 | 39.36 | 39.43 | 0.13 | 28.16 | 27.66 | 0.23 |
| 3 | 8 | 55.46 | 55.27 | 0.24 | 31.95 | 28.30 | 1.05 |
| 3 | 10 | 70.23 | 69.89 | 0.33 | 34.88 | 28.48 | 1.96 |
| 5 | 5 | 46.26 | 46.39 | 0.18 | 29.37 | 28.72 | 0.27 |
| 5 | 8 | 72.98 | 72.87 | 0.27 | 34.53 | 29.97 | 1.18 |
| 5 | 10 | 97.50 | 97.25 | 0.33 | 38.31 | 30.33 | 2.25 |
| 7 | 5 | 53.12 | 53.34 | 0.24 | 31.35 | 30.59 | 0.27 |
| 7 | 8 | 90.42 | 90.44 | 0.31 | 37.99 | 32.92 | 1.07 |
| 7 | 10 | 124.68 | 124.55 | 0.34 | 42.59 | 33.57 | 2.07 |
| 10 | 5 | 63.30 | 63.65 | 0.31 | 32.53 | 32.41 | 0.25 |
| 10 | 8 | 116.40 | 116.67 | 0.55 | 40.39 | 37.40 | 0.73 |
| 10 | 10 | 165.22 | 165.30 | 0.58 | 45.71 | 39.04 | 1.28 |

% `cont` stands for the percentage of contamination; `sdt` stands for the number of standard deviations of the outliers.

Table 2S. The overall mean value (om) of the estimated genotypic means ($\widehat{\boldsymbol{\mu}}$) and breeding values ($\widehat{\mathbf{g}}$) together with the corresponding Pearson correlation coefficients ($r_s$) between the estimates of $\widehat{\boldsymbol{\mu}}$ and $\widehat{\mathbf{g}}$ and the true breeding values, obtained using the classical (CLS) and the robust (ROB) methods under the **random** contamination scenarios. The true overall genotypic mean is 8.923 (computed as the average of the true 698 breeding values $\mathbf{g}$).

| **Scenarios I** | | **1st-stage ($\widehat{\boldsymbol{\mu}}$)** | | **2nd-stage ($\widehat{\mathbf{g}}$)** | |
|---|---|---|---|---|---|
| % cont | sdt | **CLS** (om/$r_s$) | **ROB** (om/$r_s$) | **CLS** (om/$r_s$) | **ROB** (om/$r_s$) |
| 0 | - | 8.908/0.76 | 8.906/0.76 | 5.001/0.90 | 4.935/0.90 |
| 1 | 5 | 9.092/0.75 | 9.090/0.75 | 4.925/0.90 | 4.880/0.90 |
| 1 | 8 | 9.203/0.72 | 9.200/0.72 | 4.823/0.89 | 4.861/0.90 |
| 1 | 10 | 9.276/0.70 | 9.274/0.70 | 4.737/0.88 | 4.856/0.90 |
| 3 | 5 | 9.460/0.71 | 9.458/0.71 | 4.822/0.89 | 4.791/0.89 |
| 3 | 8 | 9.792/0.65 | 9.789/0.65 | 4.586/0.86 | 4.729/0.89 |
| 3 | 10 | 10.013/0.61 | 10.010/0.61 | 4.405/0.85 | 4.713/0.89 |
| 5 | 5 | 9.829/0.69 | 9.827/0.69 | 4.758/0.88 | 4.730/0.89 |
| 5 | 8 | 10.381/0.60 | 10.379/0.60 | 4.442/0.85 | 4.612/0.88 |
| 5 | 10 | 10.749/0.55 | 10.747/0.55 | 4.214/0.82 | 4.577/0.88 |
| 7 | 5 | 10.196/0.67 | 10.194/0.66 | 4.599/0.87 | 4.576/0.88 |
| 7 | 8 | 10.969/0.57 | 10.967/0.57 | 4.189/0.83 | 4.374/0.87 |
| 7 | 10 | 11.484/0.51 | 11.483/0.51 | 3.918/0.80 | 4.315/0.87 |
| 10 | 5 | 10.748/0.64 | 10.748/0.63 | 4.562/0.86 | 4.472/0.87 |
| 10 | 8 | 11.853/0.52 | 11.853/0.52 | 4.092/0.81 | 4.0787/0.85 |
| 10 | 10 | 12.589/0.46 | 12.589/0.46 | 3.789/0.78 | 3.948/0.84 |

% `cont` stands for the percentage of contamination; `sdt` stands for the number of standard deviations of the outliers.

Table 3S. Mean squared deviations of the estimated genotypic means from the true breeding values – $\text{MSD}_\mu = \sum_{i=1}^{698}\sum_{j=1}^{1000}\frac{(\widehat{\mu}_{ij}-\mu_{ij})^2}{698\times 1000}$ – and mean squared deviations of the estimated breeding values from the true breeding values – $\text{MSD}_g = \sum_{i=1}^{698}\sum_{j=1}^{1000}\frac{(\widehat{g}_{ij}-g_{ij})^2}{698\times 1000}$ – for the classical (CLS) and robust (ROB) methods under the **block** contamination scenarios. MSDs of the classical from the robust estimates are also reported (CLS-ROB).

| **Scenarios II** | | **1st-stage ($\widehat{\boldsymbol{\mu}}$)** | | | **2nd-stage ($\widehat{\mathbf{g}}$)** | | |
|---|---|---|---|---|---|---|---|
| No. blocks | sdt | **CLS** | **ROB** | **CLS-ROB** | **CLS** | **ROB** | **CLS-ROB** |
| 0 | - | 28.97 | 29.08 | 0.06 | 25.18 | 25.55 | 0.05 |
| 1 | 5 | 30.90 | 29.45 | 0.91 | 25.35 | 25.61 | 0.19 |
| 1 | 8 | 30.75 | 29.69 | 1.76 | 25.38 | 25.63 | 0.27 |
| 1 | 10 | 31.16 | 29.83 | 2.31 | 25.39 | 25.63 | 0.30 |
| 2 | 5 | 30.93 | 29.53 | 1.41 | 25.32 | 25.56 | 0.21 |
| 2 | 8 | 32.49 | 29.81 | 3.08 | 25.36 | 25.58 | 0.28 |
| 2 | 10 | 33.79 | 29.93 | 4.41 | 25.37 | 25.59 | 0.31 |
| 3 | 5 | 32.06 | 29.64 | 2.06 | 25.34 | 25.65 | 0.22 |
| 3 | 8 | 35.22 | 29.96 | 5.08 | 25.37 | 25.68 | 0.29 |
| 3 | 10 | 38.02 | 30.11 | 7.74 | 25.37 | 25.67 | 0.30 |
| 4 | 5 | 33.48 | 29.56 | 2.70 | 25.34 | 25.54 | 0.18 |
| 4 | 8 | 38.74 | 29.61 | 7.25 | 25.37 | 25.54 | 0.24 |
| 4 | 10 | 43.51 | 29.63 | 11.52 | 25.37 | 25.54 | 0.25 |
| 5 | 5 | 35.31 | 29.76 | 3.69 | 25.39 | 25.58 | 0.17 |
| 5 | 8 | 43.33 | 29.83 | 10.51 | 25.40 | 25.58 | 0.22 |
| 5 | 10 | 50.66 | 29.88 | 17.03 | 25.40 | 25.59 | 0.24 |

`No Blocks` stands for the number of contaminated blocks; `sdt` stands for the number of standard deviations of the outliers.

Table 4S. The overall mean value (om) of the estimated genotypic means ($\widehat{\boldsymbol{\mu}}$) and breeding values ($\widehat{\mathbf{g}}$) together with the corresponding Pearson correlation coefficients ($r_s$) between the estimates of $\widehat{\boldsymbol{\mu}}$ and $\widehat{\mathbf{g}}$ and the true breeding values, obtained using the classical (CLS) and the robust (ROB) methods under the **block** contamination scenarios. The true overall genotypic mean is 8.923 (computed as the average of the true 698 breeding values $\mathbf{g}$).

| **Scenarios II** | | **1st-stage ($\widehat{\boldsymbol{\mu}}$)** | | **2nd-stage ($\widehat{\mathbf{g}}$)** | |
|---|---|---|---|---|---|
| No. blocks | sdt | **CLS** (om/$r_s$) | **ROB** (om/$r_s$) | **CLS** (om/$r_s$) | **ROB** (om/$r_s$) |
| 0 | - | 8.908/0.77 | 8.906/0.76 | 4.979/0.90 | 4.935/0.90 |
| 1 | 5 | 9.379/0.76 | 9.017/0.76 | 5.004/0.90 | 4.933/0.90 |
| 1 | 8 | 9.661/0.76 | 9.022/0.76 | 5.008/0.90 | 4.930/0.90 |
| 1 | 10 | 9.849/0.76 | 9.025/0.76 | 5.009/0.90 | 4.929/0.90 |
| 2 | 5 | 9.837/0.76 | 9.112/0.76 | 5.012/0.90 | 4.939/0.90 |
| 2 | 8 | 10.394/0.76 | 9.119/0.76 | 5.013/0.90 | 4.936/0.90 |
| 2 | 10 | 10.765/0.76 | 9.121/0.76 | 5.014/0.90 | 4.934/0.90 |
| 3 | 5 | 10.289/0.76 | 9.211/0.76 | 5.011/0.90 | 4.932/0.90 |
| 3 | 8 | 11.118/0.76 | 9.218/0.76 | 5.013/0.90 | 4.929/0.90 |
| 3 | 10 | 11.670/0.76 | 9.220/0.76 | 5.013/0.90 | 4.929/0.90 |
| 4 | 5 | 10.715/0.76 | 9.299/0.76 | 5.015/0.90 | 4.946/0.90 |
| 4 | 8 | 11.799/0.76 | 9.300/0.76 | 5.015/0.90 | 4.946/0.90 |
| 4 | 10 | 12.522/0.76 | 9.301/0.76 | 5.015/0.90 | 4.946/0.90 |
| 5 | 5 | 11.153/0.76 | 9.411/0.76 | 5.008/0.90 | 4.942/0.90 |
| 5 | 8 | 12.500/0.76 | 9.413/0.76 | 5.011/0.90 | 4.941/0.90 |
| 5 | 10 | 13.398/0.76 | 9.414/0.76 | 5.011/0.90 | 4.941/0.90 |

`No Blocks` stands for the number of contaminated blocks; `sdt` stands for the number of standard deviations of the outliers.

Table 5S. Mean squared deviations of the estimated breeding values and the true breeding values ($MSD_{\mathbf{g}}$) for the classical (CLS) approach computed using the Smith's and standard weights for the **block** and **random** contamination scenarios.

| **Block scenarios** | | **Smith** | **Standard** | **Random scenarios** | | **Smith** | **Standard** |
|---|---|---|---|---|---|---|---|
| No. blocks | sdt | | | % cont | sdt | | |
| 0 | - | 25.18 | 25.29 | 0 | - | 25.18 | 25.29 |
| 1 | 5 | 25.35 | 25.61 | 1 | 5 | 26.29 | 26.40 |
| 1 | 8 | 25.38 | 25.77 | 1 | 8 | 27.85 | 27.96 |
| 1 | 10 | 25.39 | 25.87 | 1 | 10 | 29.16 | 29.28 |
| 2 | 5 | 25.33 | 25.74 | 3 | 5 | 28.16 | 28.31 |
| 2 | 8 | 25.36 | 26.09 | 3 | 8 | 31.95 | 32.16 |
| 2 | 10 | 25.37 | 26.40 | 3 | 10 | 34.88 | 35.13 |
| 3 | 5 | 25.34 | 25.99 | 5 | 5 | 29.38 | 29.61 |
| 3 | 8 | 25.37 | 26.75 | 5 | 8 | 34.53 | 34.93 |
| 3 | 10 | 25.37 | 27.49 | 5 | 10 | 38.31 | 38.85 |
| 4 | 5 | 25.34 | 26.31 | 7 | 5 | 31.35 | 31.70 |
| 4 | 8 | 25.37 | 27.70 | 7 | 8 | 37.99 | 38.64 |
| 4 | 10 | 25.37 | 29.08 | 7 | 10 | 42.59 | 43.48 |
| 5 | 5 | 25.39 | 26.80 | 10 | 5 | 32.53 | 33.20 |
| 5 | 8 | 25.40 | 29.05 | 10 | 8 | 40.39 | 41.73 |
| 5 | 10 | 25.40 | 31.27 | 10 | 10 | 45.71 | 47.56 |

`% cont` stands for percentage of contamination; `sdt` stands for the number of standard deviations of the outliers;

`No Blocks` stands for the number of contaminated blocks.

Table 6S. The mean squared deviation of heritability – $\mathrm{MSD}_{\mathtt{H}} = \sum_{j=1}^{1000} \frac{(\widehat{H}_j^2 - (r_{g,\widehat{g},j})^2)^2}{1000}$ (Method 5) – and predictive accuracy – $\mathrm{MSD}_{\mathtt{PA}} = \sum_{j=1}^{1000} \frac{(\widehat{r}_{g,\widehat{g},j} - r_{g,\widehat{g},j})^2}{1000}$ (Methods 5 and 7) – of the classical (CLS) from the robust (ROB) method under the **random** contamination scenarios.

| **Random scenarios** | | H | | PA – **M5** | | PA – **M7** | |
|---|---|---|---|---|---|---|---|
| % cont | sdt | **CLS** | **ROB** | **CLS** | **ROB** | **CLS** | **ROB** |
| 0 | - | 0.01 | 0.01 | 0.00 | 0.00 | 0.01 | 0.01 |
| 1 | 5 | 0.02 | 0.01 | 0.00 | 0.00 | 0.01 | 0.01 |
| 1 | 8 | 0.02 | 0.01 | 0.00 | 0.00 | 0.01 | 0.01 |
| 1 | 10 | 0.02 | 0.01 | 0.00 | 0.00 | 0.01 | 0.01 |
| 3 | 5 | 0.02 | 0.02 | 0.00 | 0.00 | 0.01 | 0.01 |
| 3 | 8 | 0.03 | 0.02 | 0.00 | 0.00 | 0.01 | 0.01 |
| 3 | 10 | 0.03 | 0.02 | 0.00 | 0.00 | 0.02 | 0.01 |
| 5 | 5 | 0.02 | 0.02 | 0.00 | 0.00 | 0.01 | 0.01 |
| 5 | 8 | 0.04 | 0.02 | 0.00 | 0.00 | 0.02 | 0.02 |
| 5 | 10 | 0.06 | 0.02 | 0.01 | 0.00 | 0.02 | 0.02 |
| 7 | 5 | 0.03 | 0.03 | 0.00 | 0.00 | 0.01 | 0.02 |
| 7 | 8 | 0.05 | 0.03 | 0.01 | 0.00 | 0.02 | 0.03 |
| 7 | 10 | 0.07 | 0.04 | 0.01 | 0.00 | 0.03 | 0.03 |
| 10 | 5 | 0.03 | 0.03 | 0.00 | 0.00 | 0.02 | 0.03 |
| 10 | 8 | 0.07 | 0.06 | 0.01 | 0.01 | 0.03 | 0.06 |
| 10 | 10 | 0.09 | 0.07 | 0.02 | 0.01 | 0.04 | 0.07 |

% cont stands for percentage of contamination; sdt stands for the number of standard deviations of the outliers.

Table 7S. Summary table of comparative perfomance of the competing methods for the block and random contamination scenarios; performance for the NULL scenarios was similar and is thus not shown.

| **Method** | **Scenario** | **1st-stage** | | | | | **2st-stage** | | | **3rd-stage** | |
|---|---|---|---|---|---|---|---|---|---|---|---|
| | | **MSD**$_\mu$ | r$_s$ | $\sigma^2_r$ | $\sigma^2_{r:b}$ | $\sigma^2_e$ | **MSD**$_g$ | r$_s$ | $\sigma^2_g$ | H2 | PA |
| CLS | random | $s$ | $s$ | $w$ | $b$ | $w$ | $w$ | $w$ | $w$ | $w$ | $w$ |
| | block | $w$ | $s$ | $w$ | $w$ | $s$ | $s$ | $s$ | $w$ | $b$ | $b$ |
| ROB | random | $s$ | $s$ | $b$ | $w$ | $b$ | $b$ | $b$ | $b$ | $b$ | $b$ |
| | bock | $b$ | $s$ | $b$ | $b$ | $s$ | $s$ | $s$ | $b$ | $w$ | $w$ |

"$b$" means *better* and "$w$" means *worse* performance than the competing method;

"$s$" means *similar* performance as the competing method.

### Appendix B

#### Supplementary figures

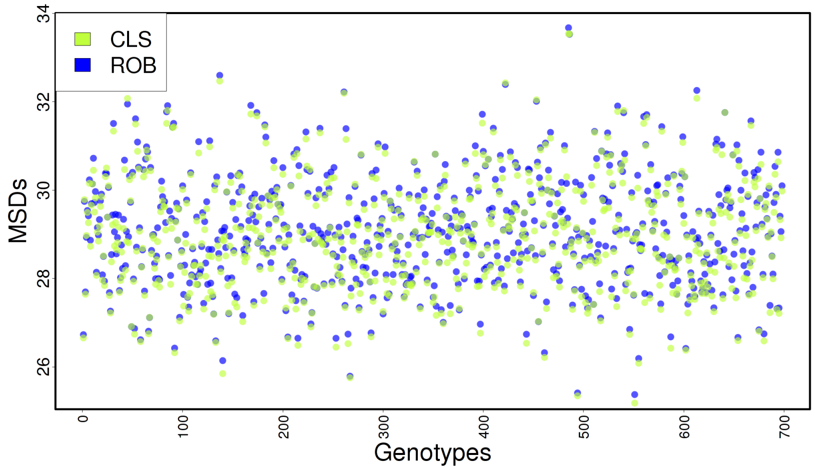


Figure 1S. Plot of the classical and robust $\text{MSD}_{\mu}^{i} = \sum_{j=1}^{1000} \frac{(\widehat{\mu}_{ij} - \mu_{ij})^2}{1000}$ for each of the 698 genotypes for the **null** scenario (`1st stage`)

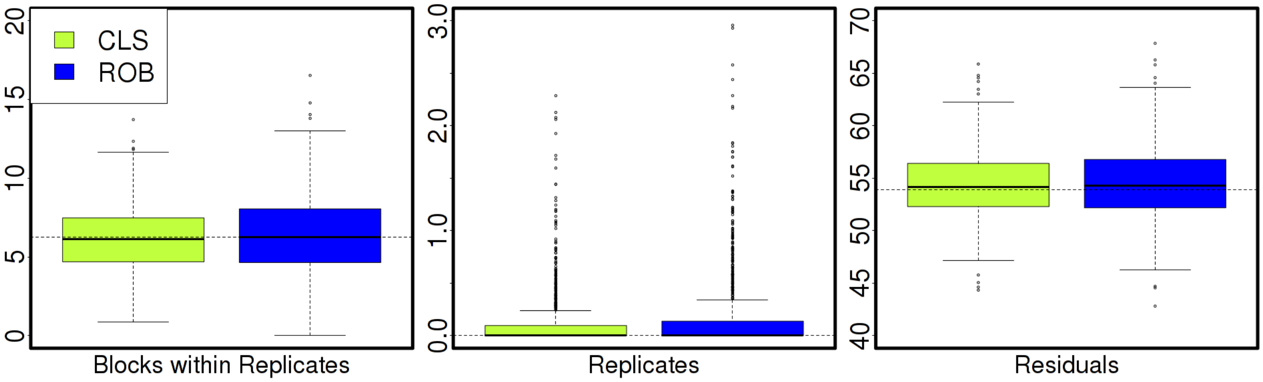


Figure 2S. Boxplots of the 1000 classical and robust estimated *block* ($\sigma^2_{r:b}$), *replicate* ($\sigma^2_r$) and *residual* variances ($\sigma^2_e$) for the **null** scenario (`1st stage`)

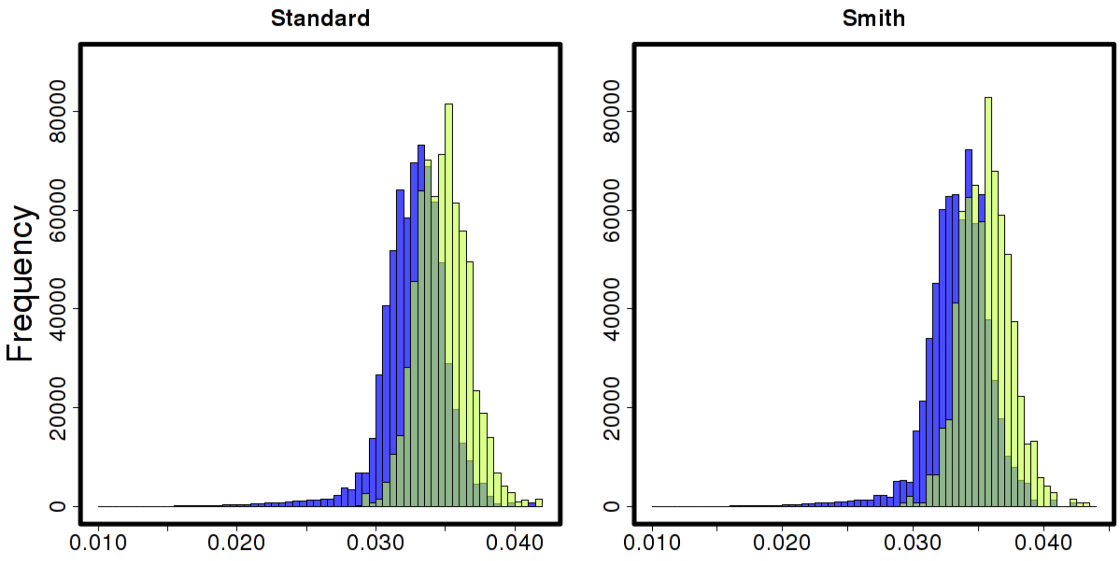


Figure 3S: Histograms of the classical and robust Standard and Smith's weights for the **null** scenario (`1st stage`)

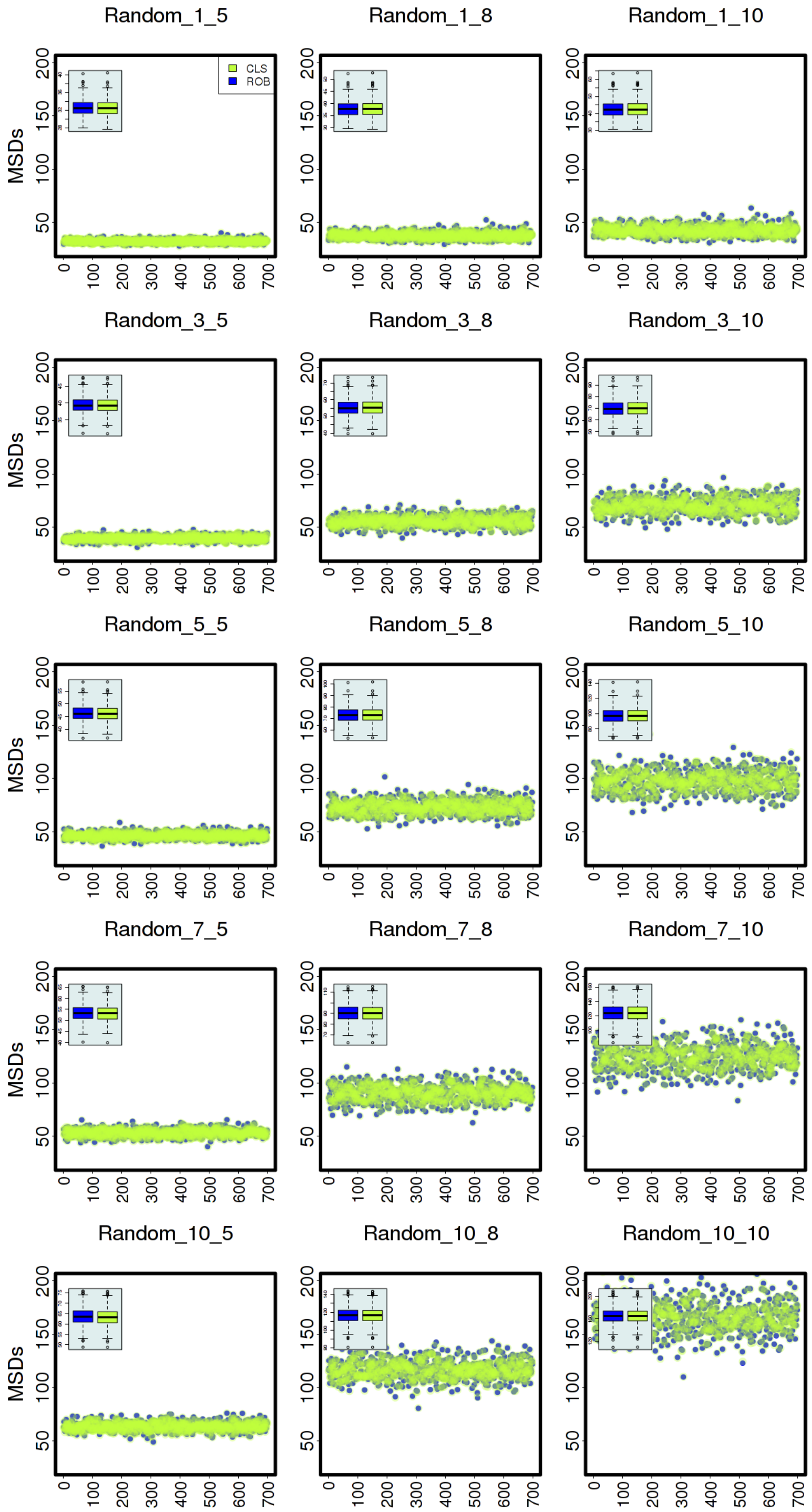


Figure 4S. Plots of classical and robust $\text{MSD}^i_\mu = \sum_{j=1}^{1000} \frac{(\widehat{\mu}_{ij} - \mu_{ij})^2}{1000}$ for each of the 698 genotypes for the **random** contamination scenarios (`1st stage`)

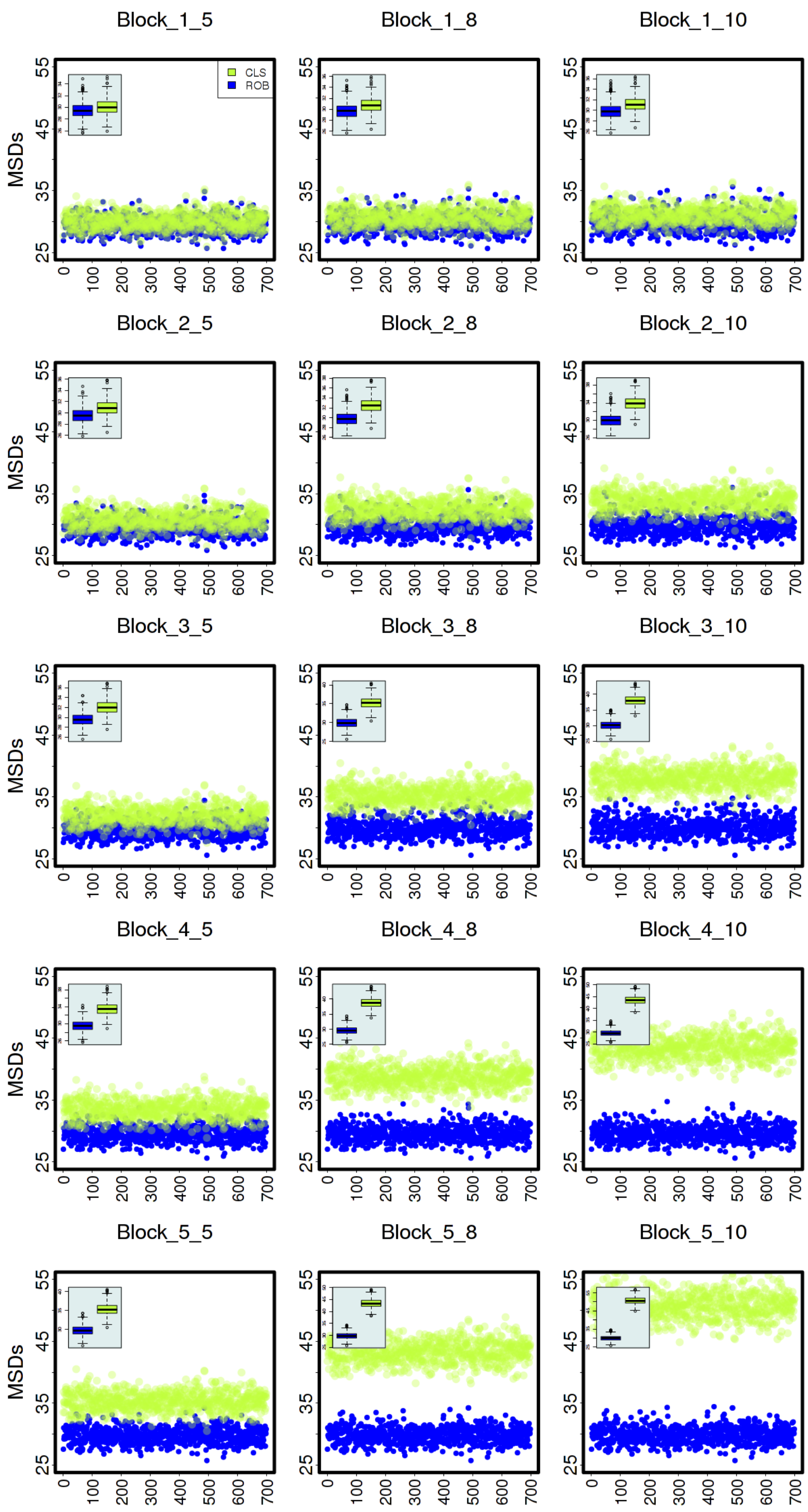


Figure 5S. Plots of classical and robust $\mathrm{MSD}_{\mu}^{i} = \sum_{j=1}^{1000} \frac{(\widehat{\mu}_{ij} - \mu_{ij})^2}{1000}$ for each of the 698 genotypes for the **block** contamination scenarios (`1st stage`)

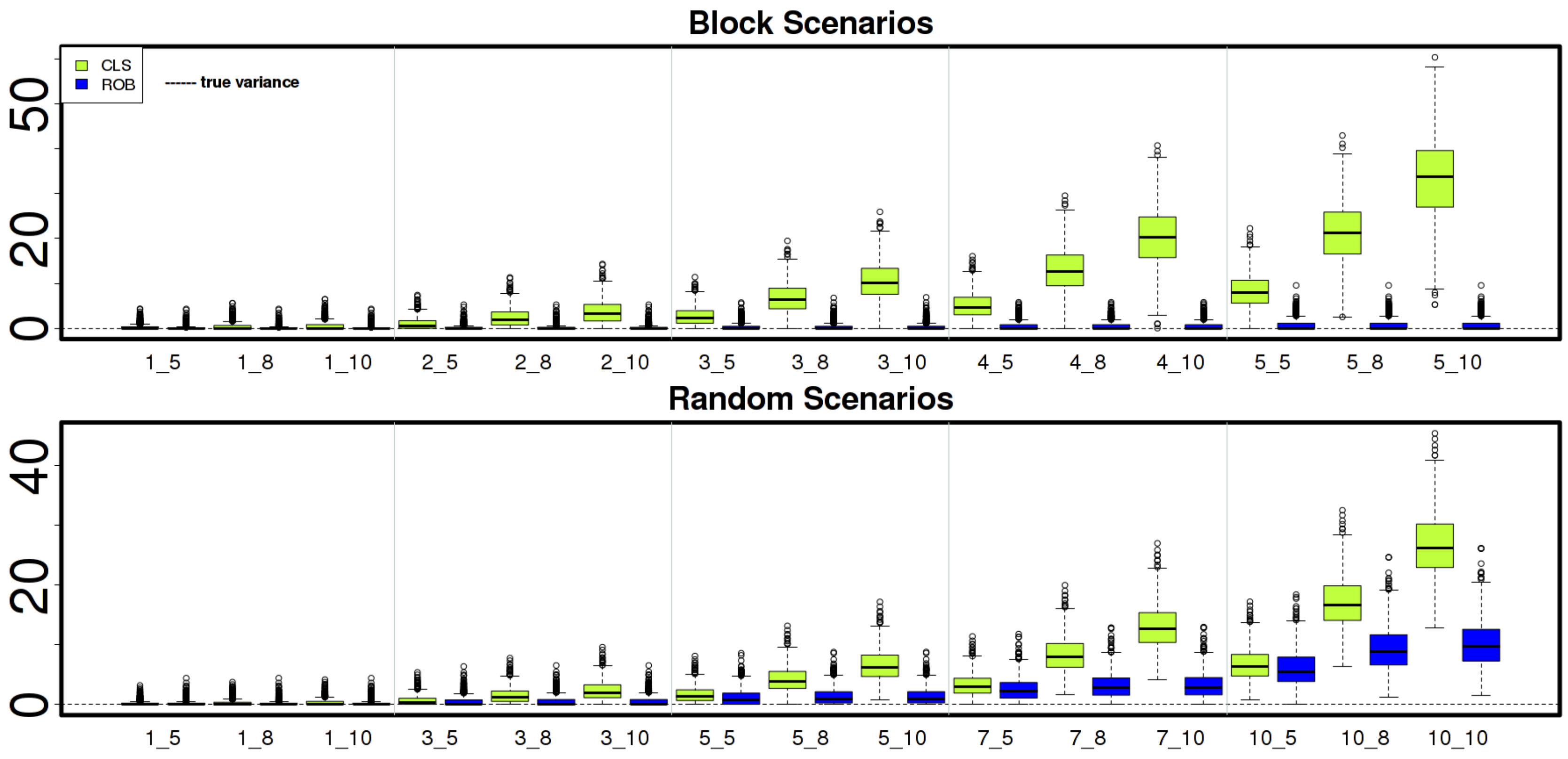


Figure 6S. Boxplots of the 1000 classical and robust estimated variances for *replicates* ($\sigma_r^2$) for the **block** and **random** contamination scenarios (`1st stage`)

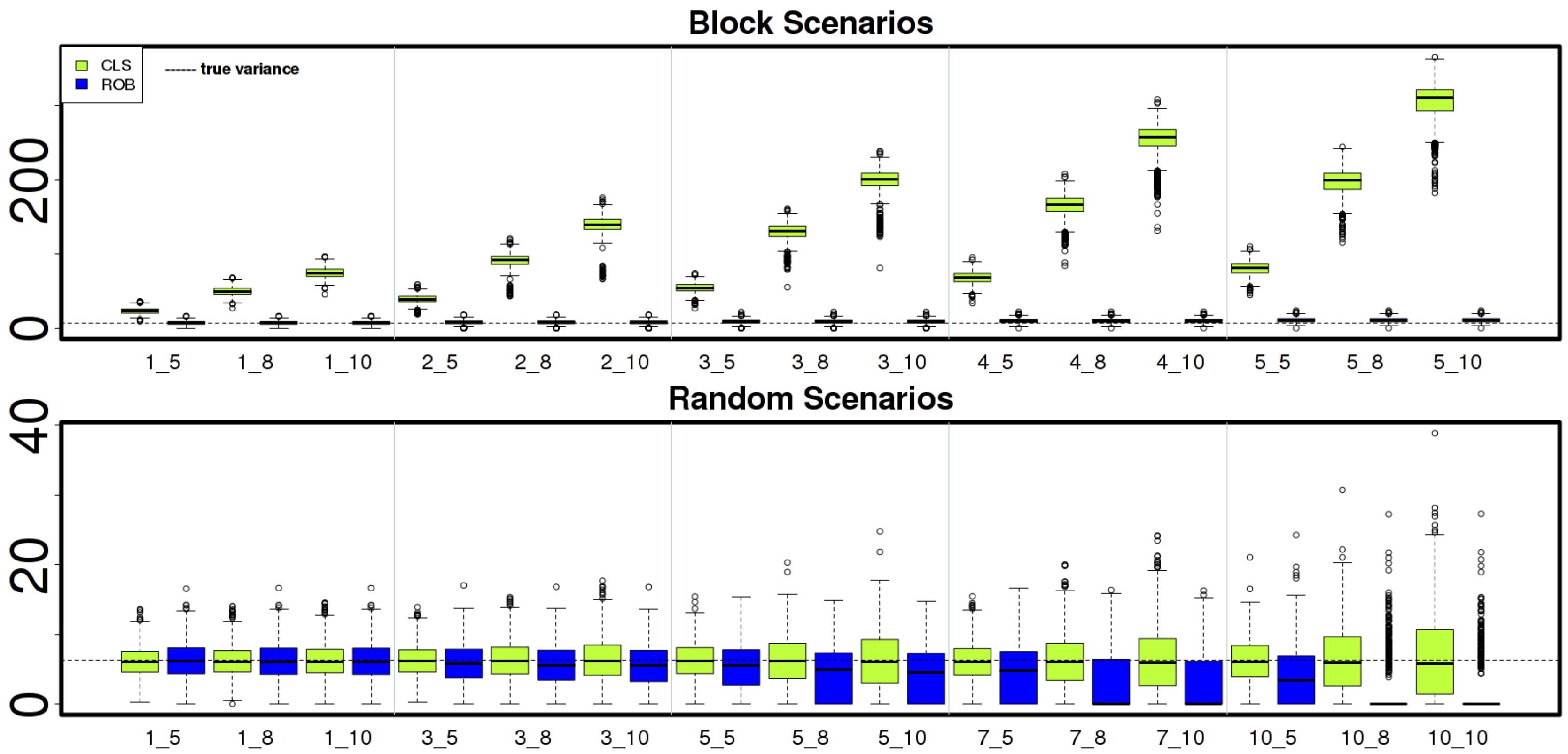


Figure 7S: Boxplots of the 1000 classical and robust estimated variances for *block nested within replicates* ($\sigma^2_{r:b}$) for the **block** and **random** contamination scenarios (`1st stage`)

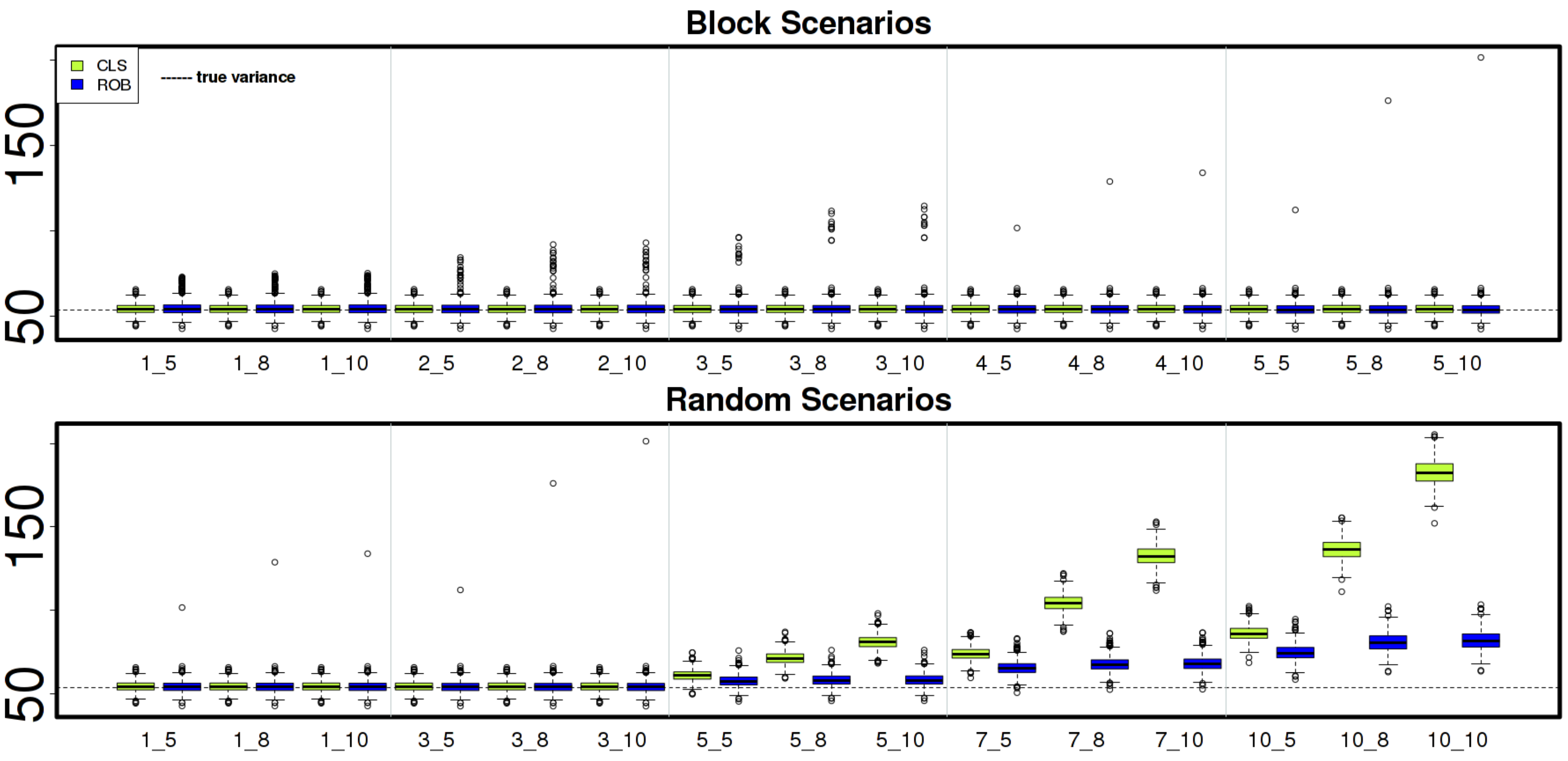


Figure 8S: Boxplots of the 1000 classical and robust estimated variances for *residuals* ($\sigma_e^2$) for the **block** and **random** contamination scenarios (`1st stage`)

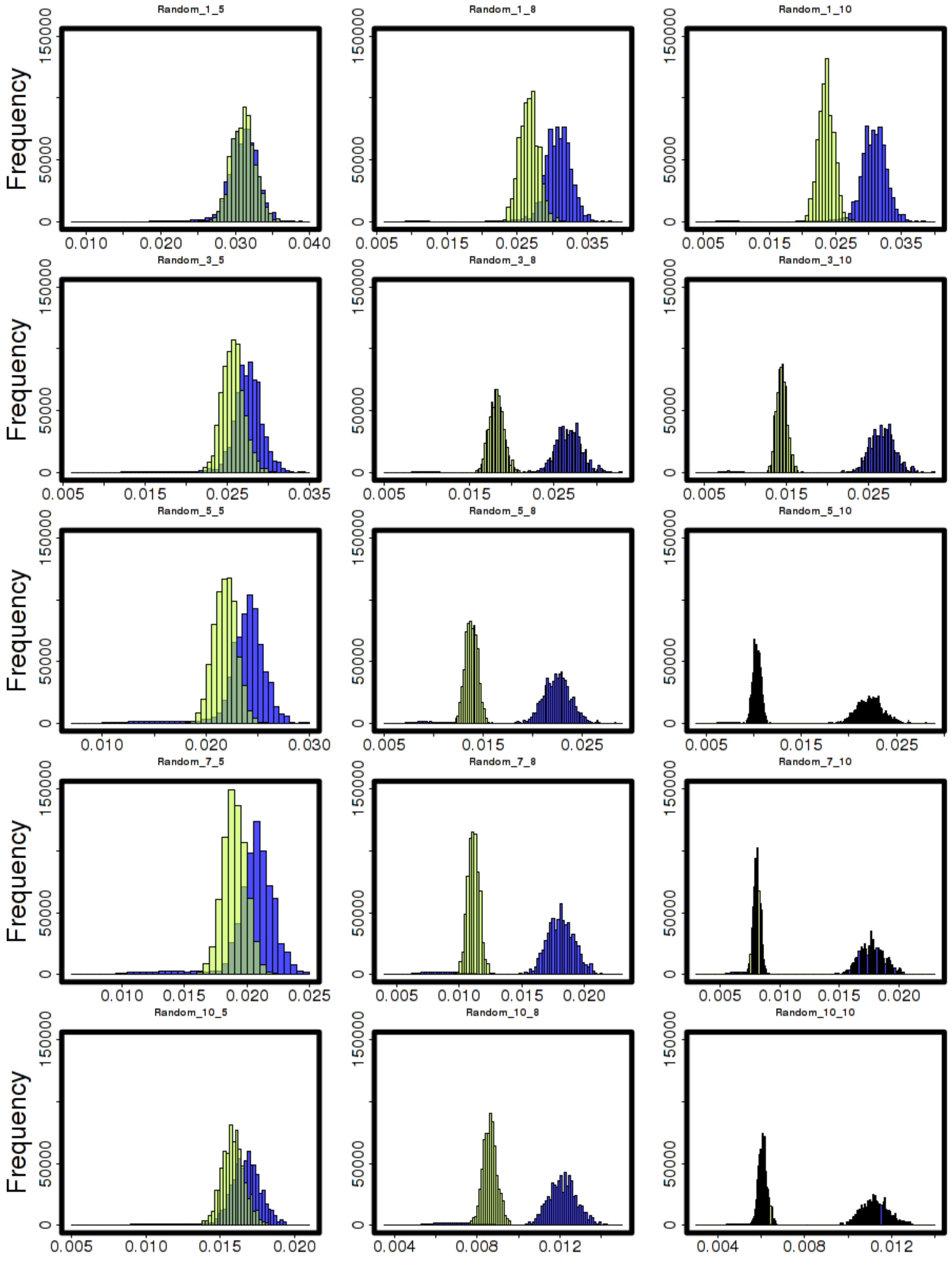


Figure 9S. Histograms of the classical and robust Standard's weights for the **random** contamination scenarios (`1st stage`)

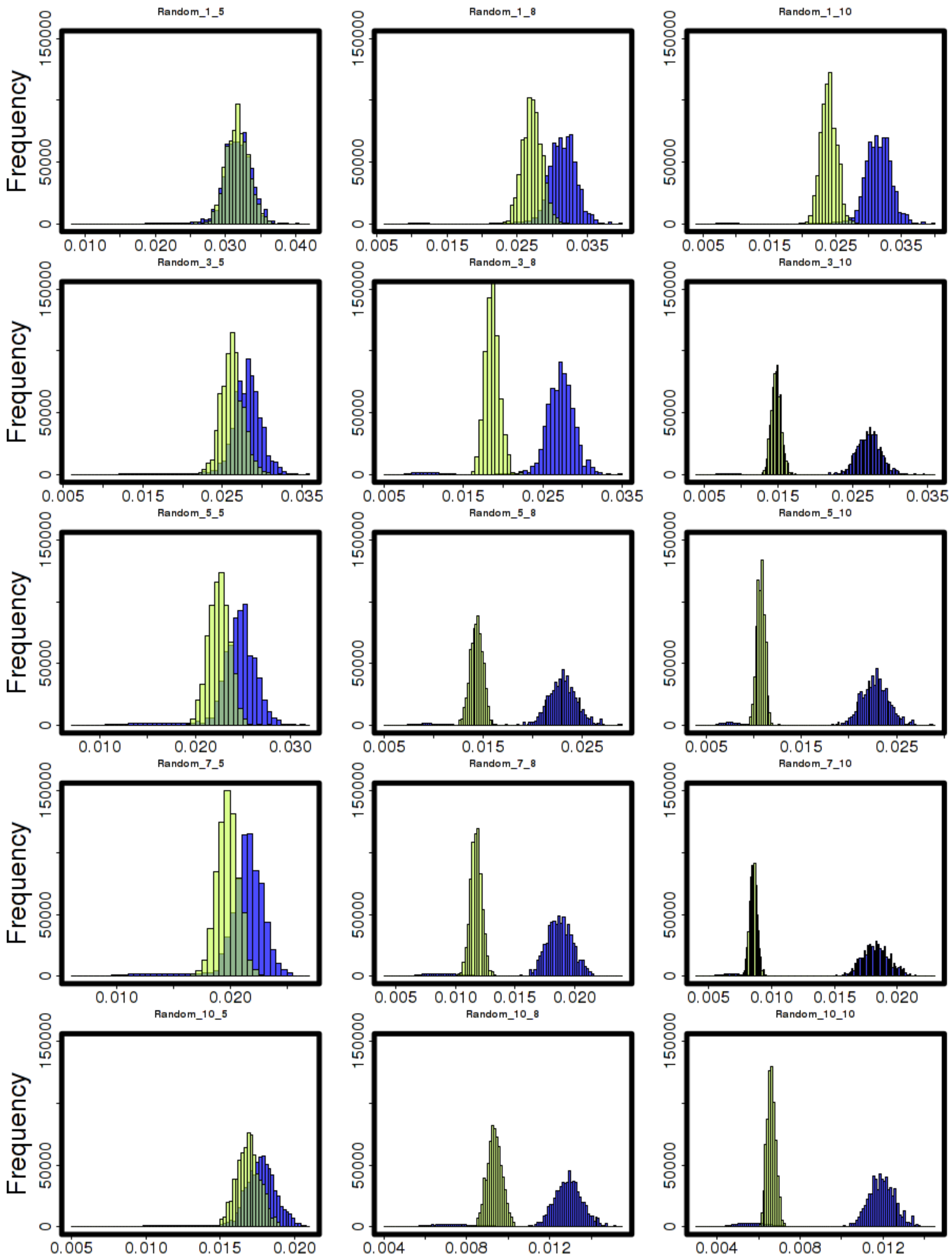


Figure 10S. Histograms of the classical and robust Smith's weights for the **random** contamination scenarios (`1st stage`)

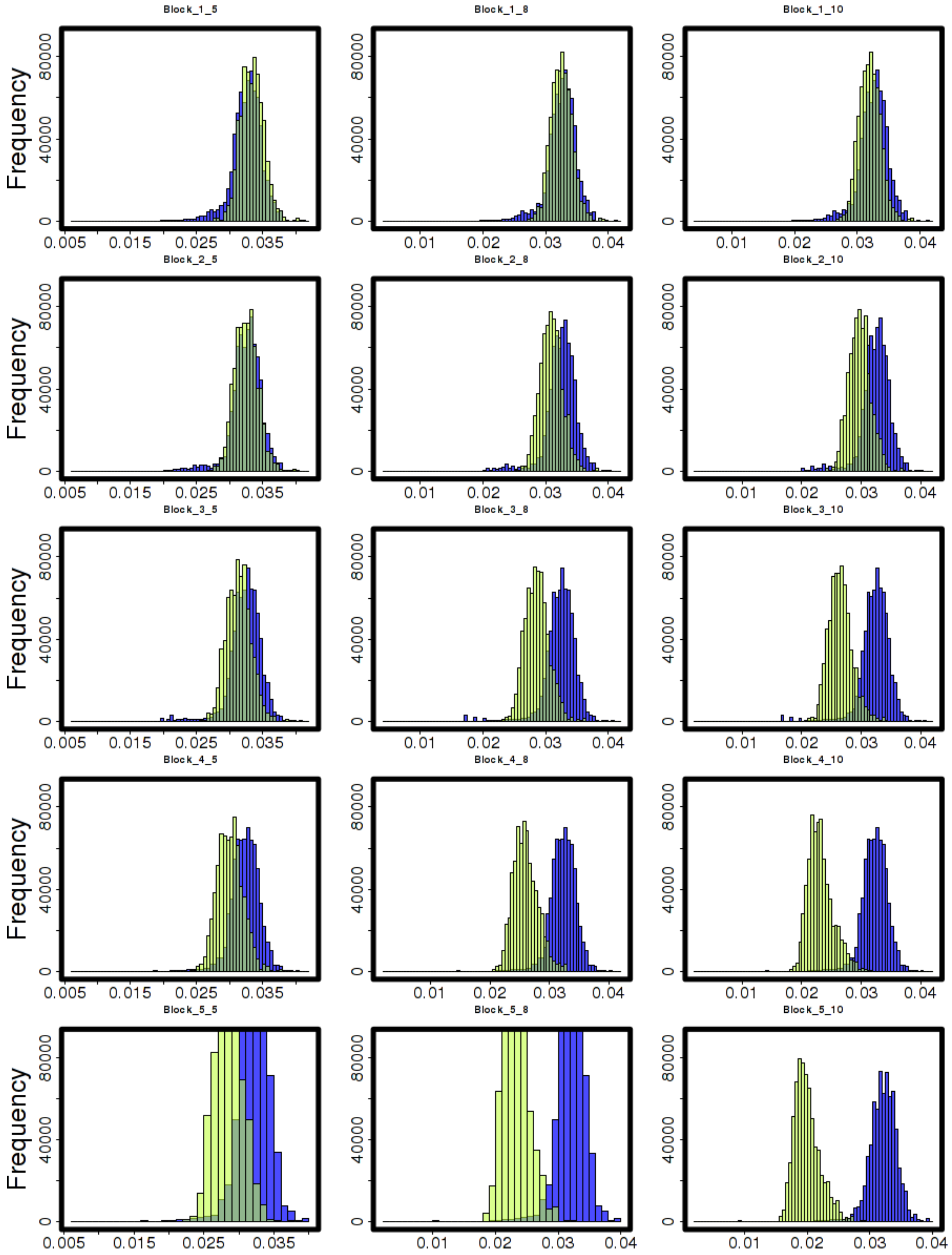


Figure 11S. Histograms of the classical and robust Standard's weights for the **block** contamination scenarios (`1st stage`)

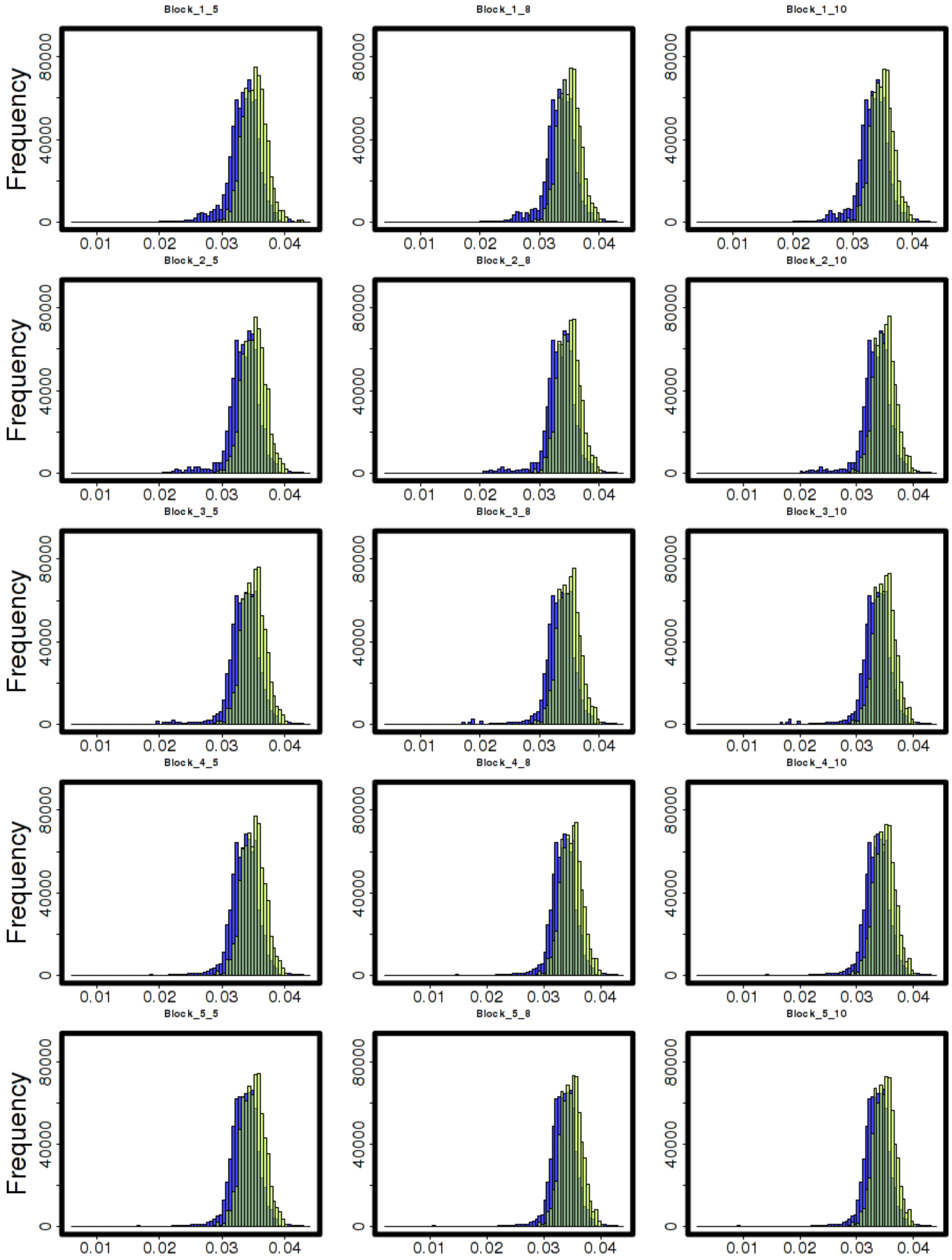


Figure 12S. Histograms of the classical and robust Smith's weights for the **block** contamination scenarios (`1st stage`)

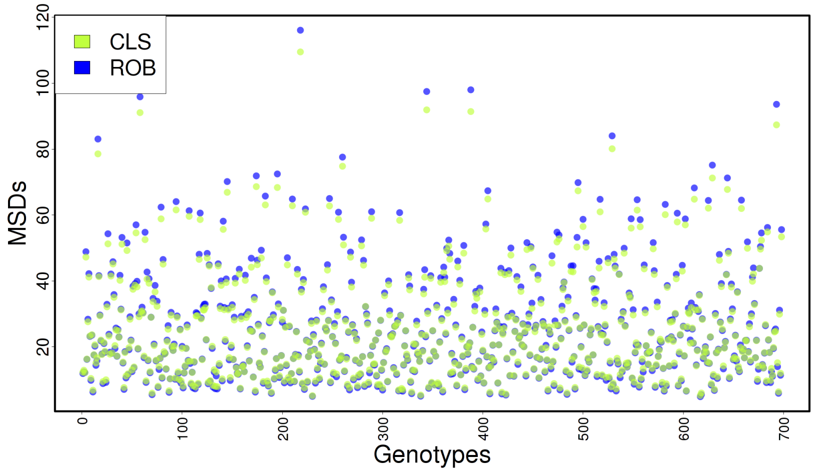


Figure 13S. Plot of the classical and robust $\text{MSD}_g^i = \sum_{j=1}^{1000} \frac{(\widehat{g}_{ij} - g_{ij})^2}{1000}$ for each of the 698 genotypes for the **null** scenario (`2nd stage`)

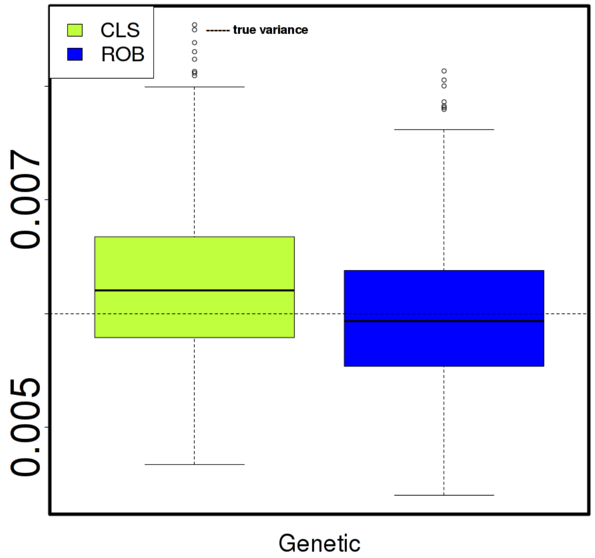


Figure 14S. Boxplots of the 1000 classical and robust estimated *genetic* ($\sigma_u^2$) variances for the **null** scenario (`2nd stage`)

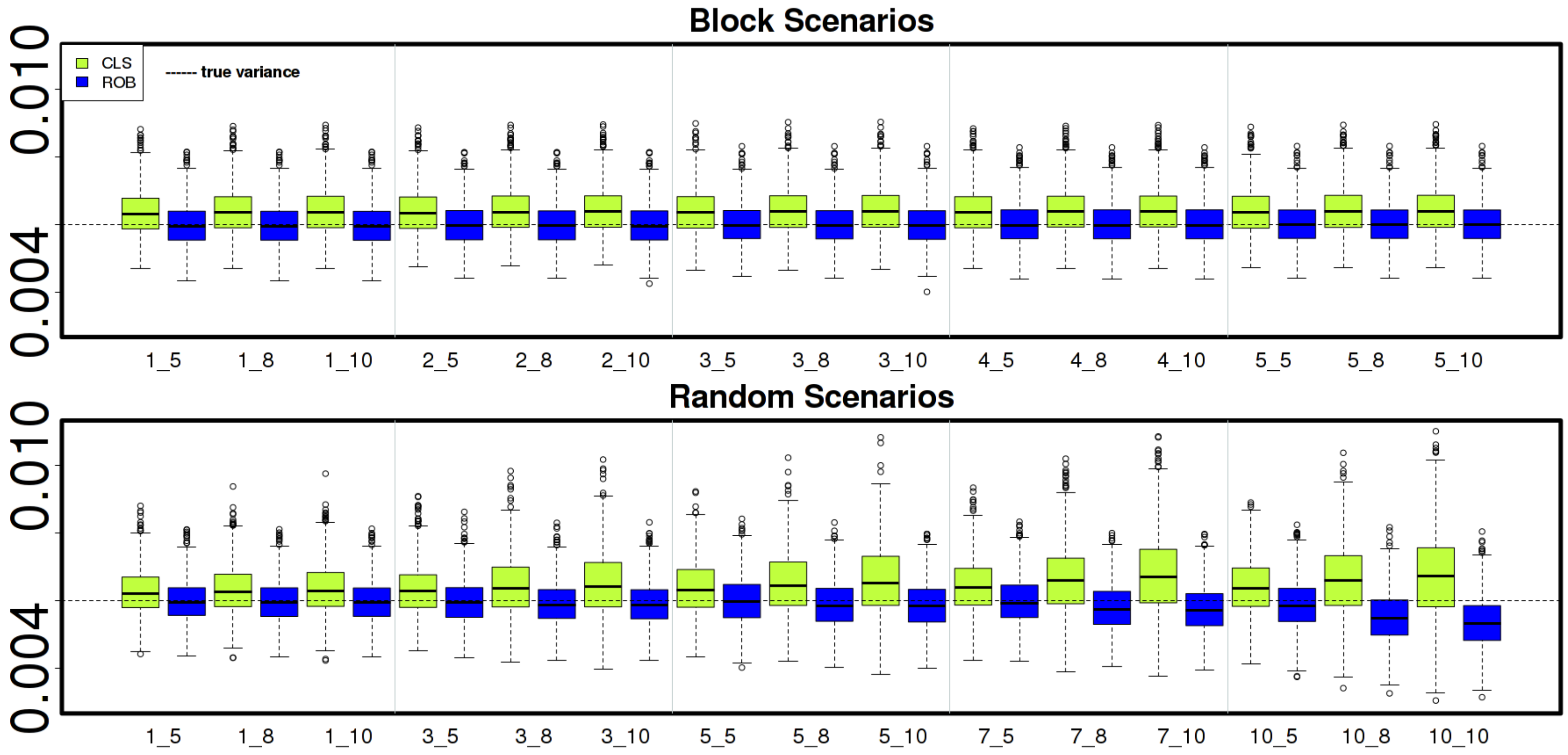


Figure 15S: Boxplots of the 1000 classical and robust estimated *genetic* variances ($\sigma_u^2$) for the **block** and **random** contamination scenarios (`2nd stage`)

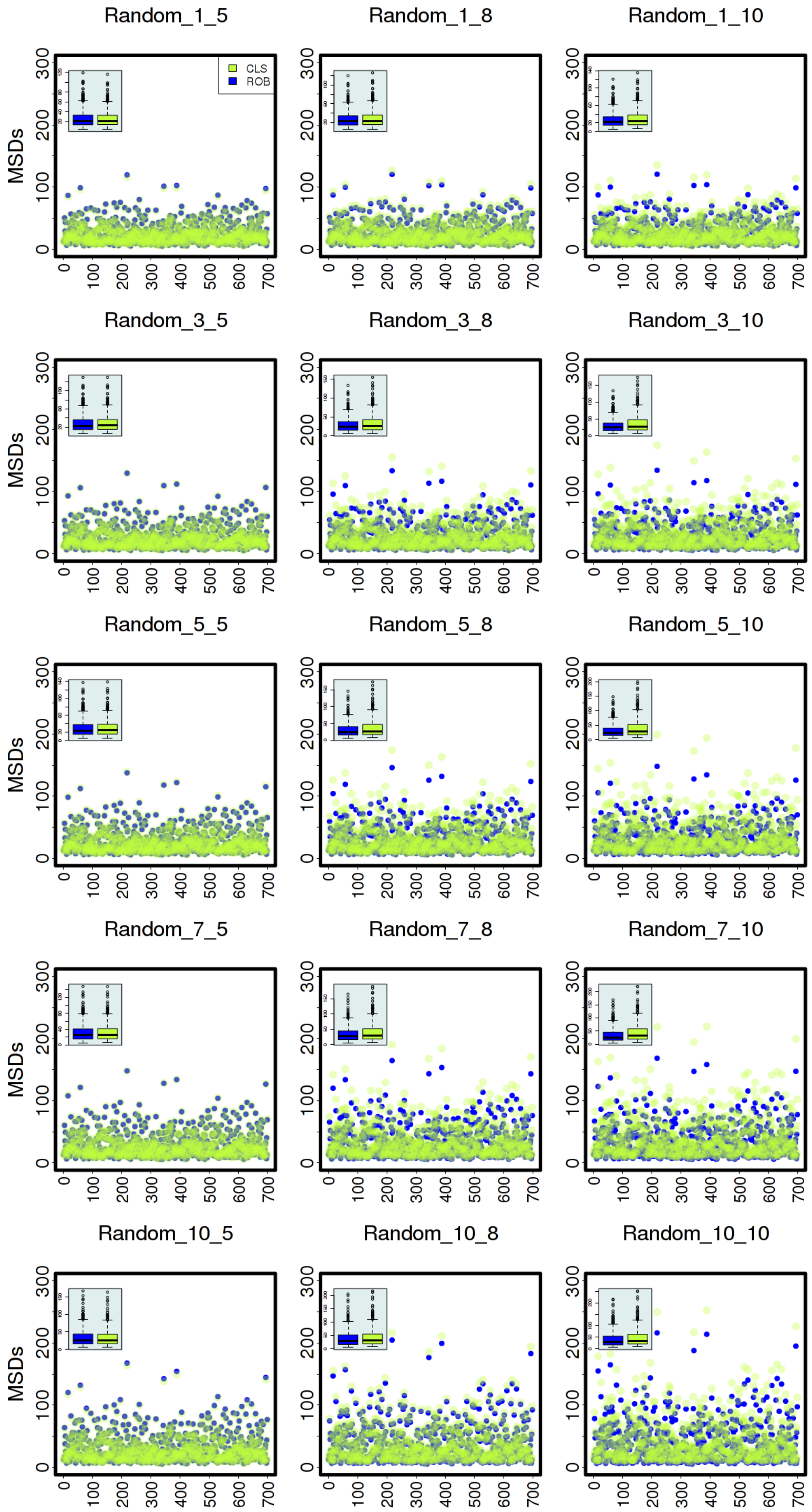


Figure 16S. Plots of the classical and robust $\mathrm{MSD}_g^i = \sum_{j=1}^{1000} \frac{(\widehat{g}_{ij} - g_{ij})^2}{1000}$ for each of the 698 genotypes for the **random** contamination scenarios (`2nd stage`)

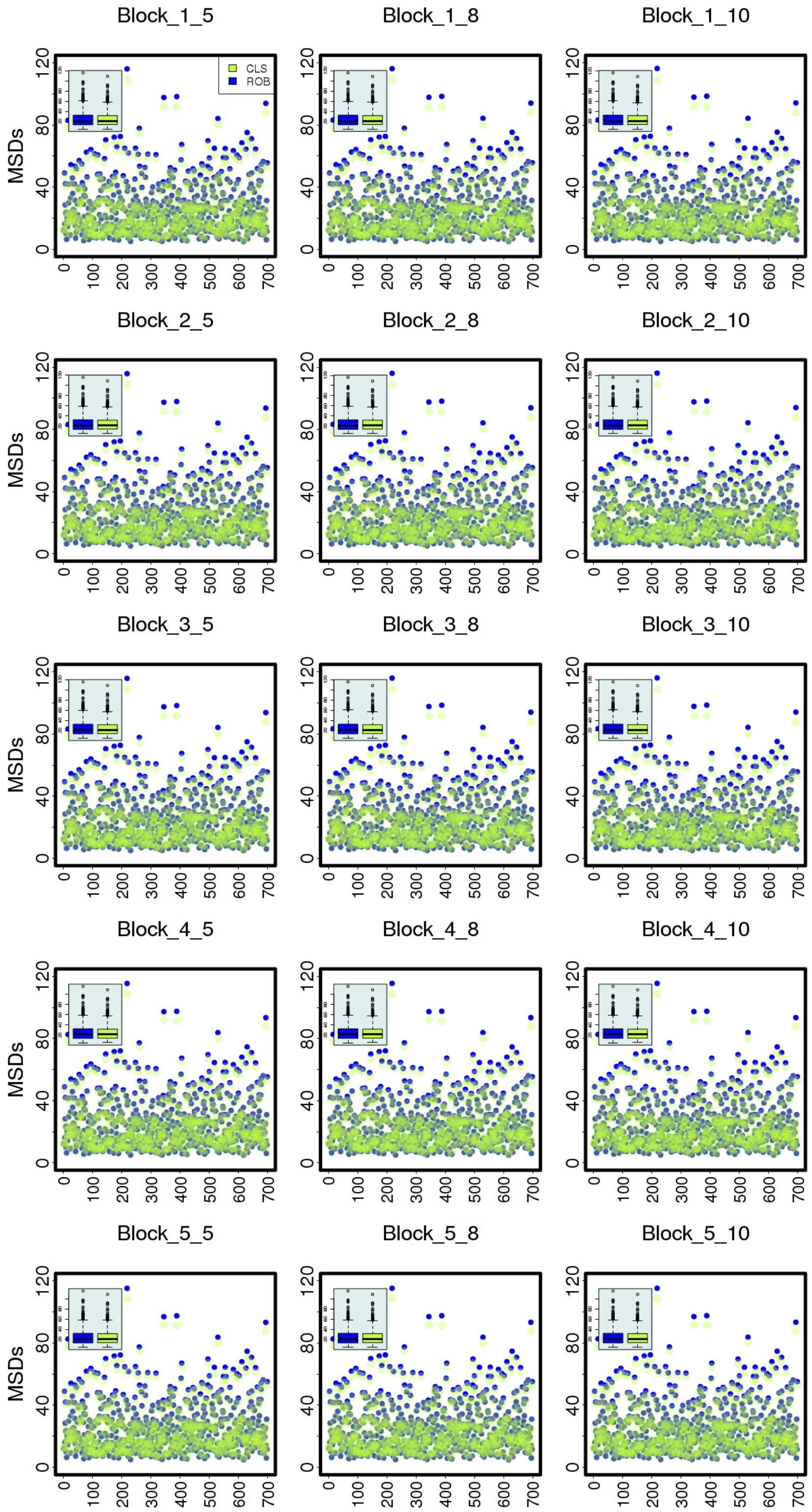


Figure 17S. Plots of the classical and robust $\mathrm{MSD}_g^i = \sum_{j=1}^{1000} \frac{(\widehat{g}_{ij} - g_{ij})^2}{1000}$ for each of the 698 genotypes for the **block** contamination scenarios (`2nd stage`)

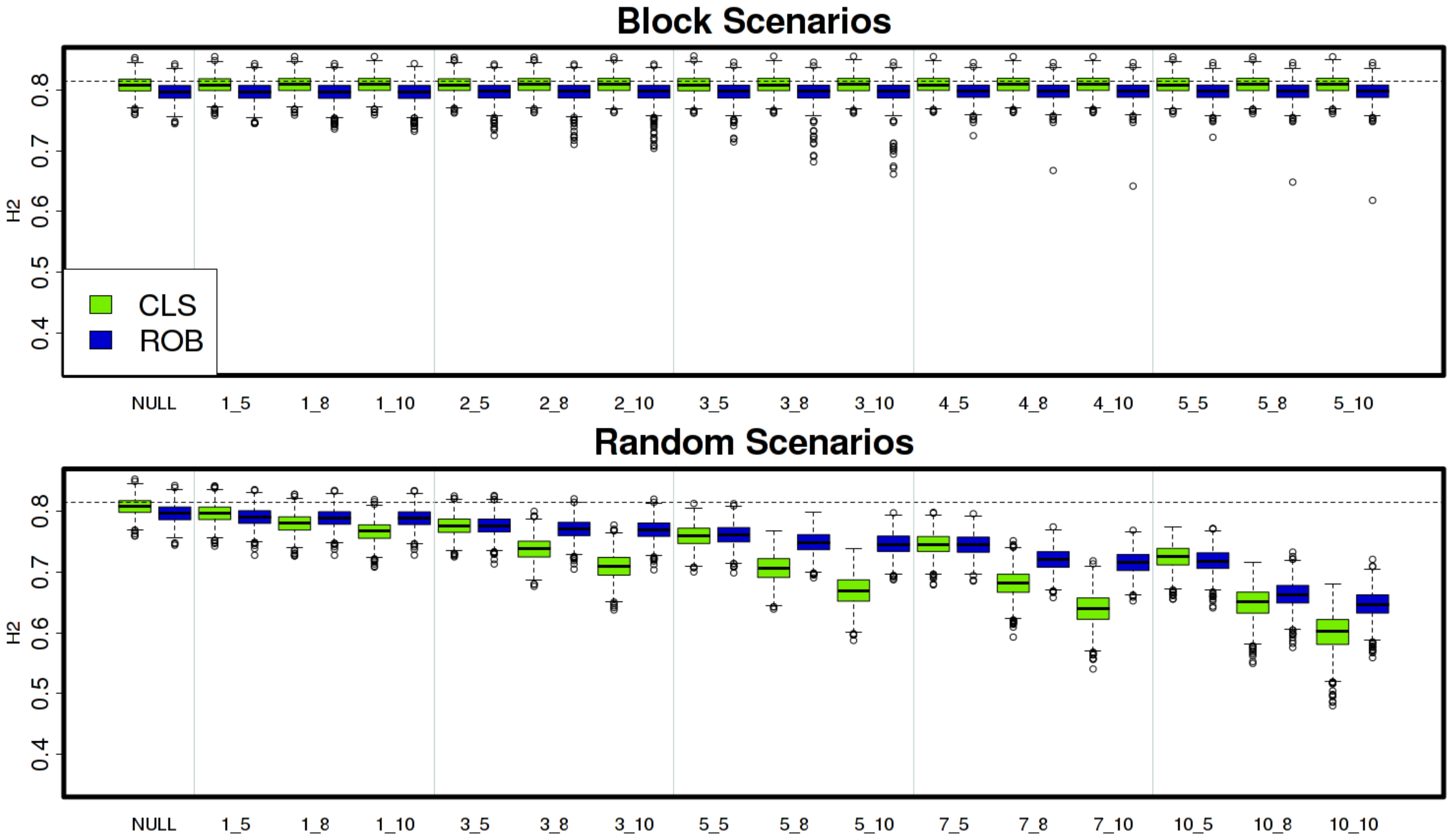


Figure 18S. Boxplots of estimated heritabilities ($H^2$) computed by method **M5** for the **block** and **random** contamination scenarios (`3rd stage`)

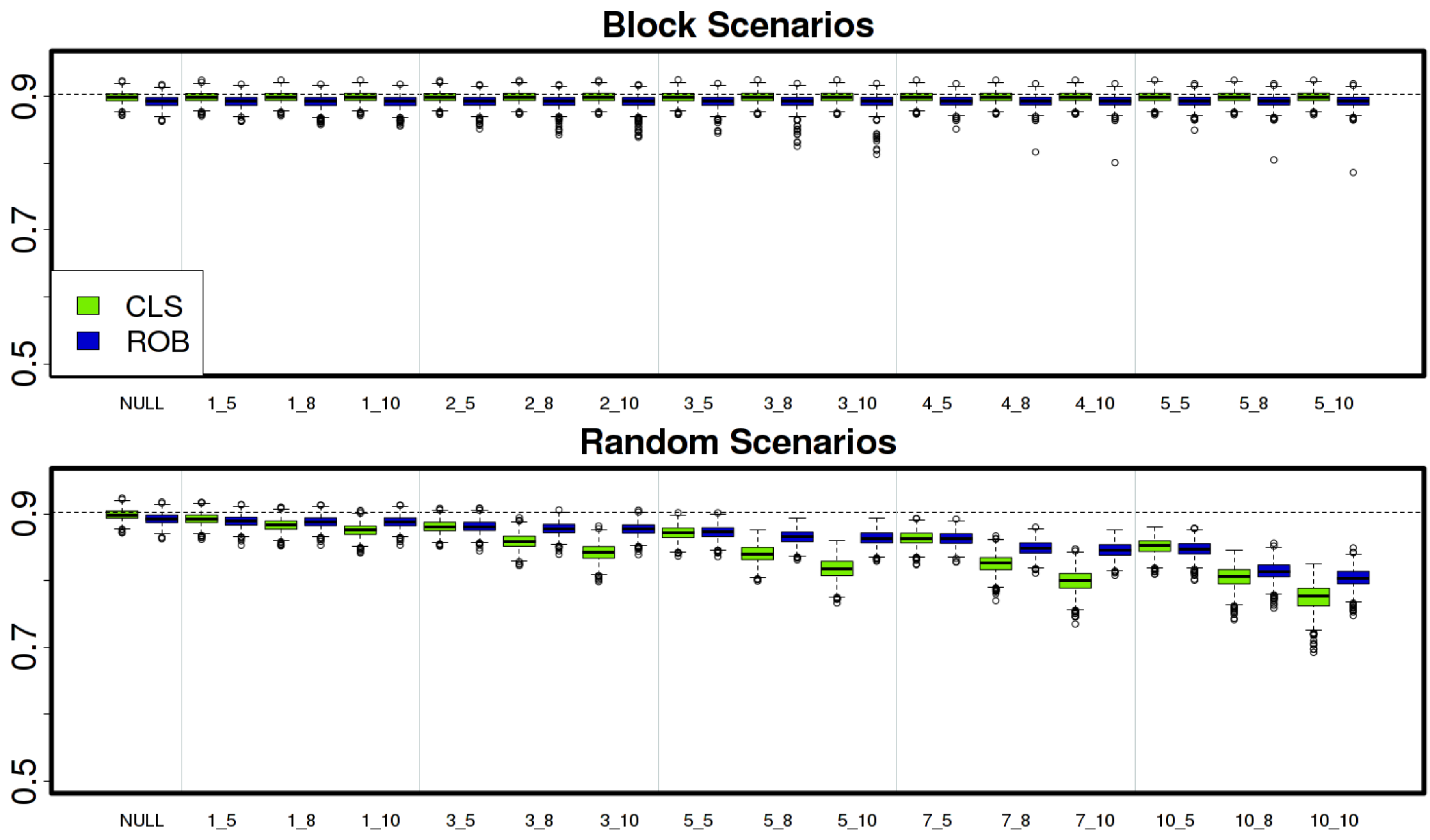


Figure 19S. Boxplots of estimated predictive accuracies (PA) computed by method **M5** for the **block** and **random** contamination scenarios (`3rd stage`)

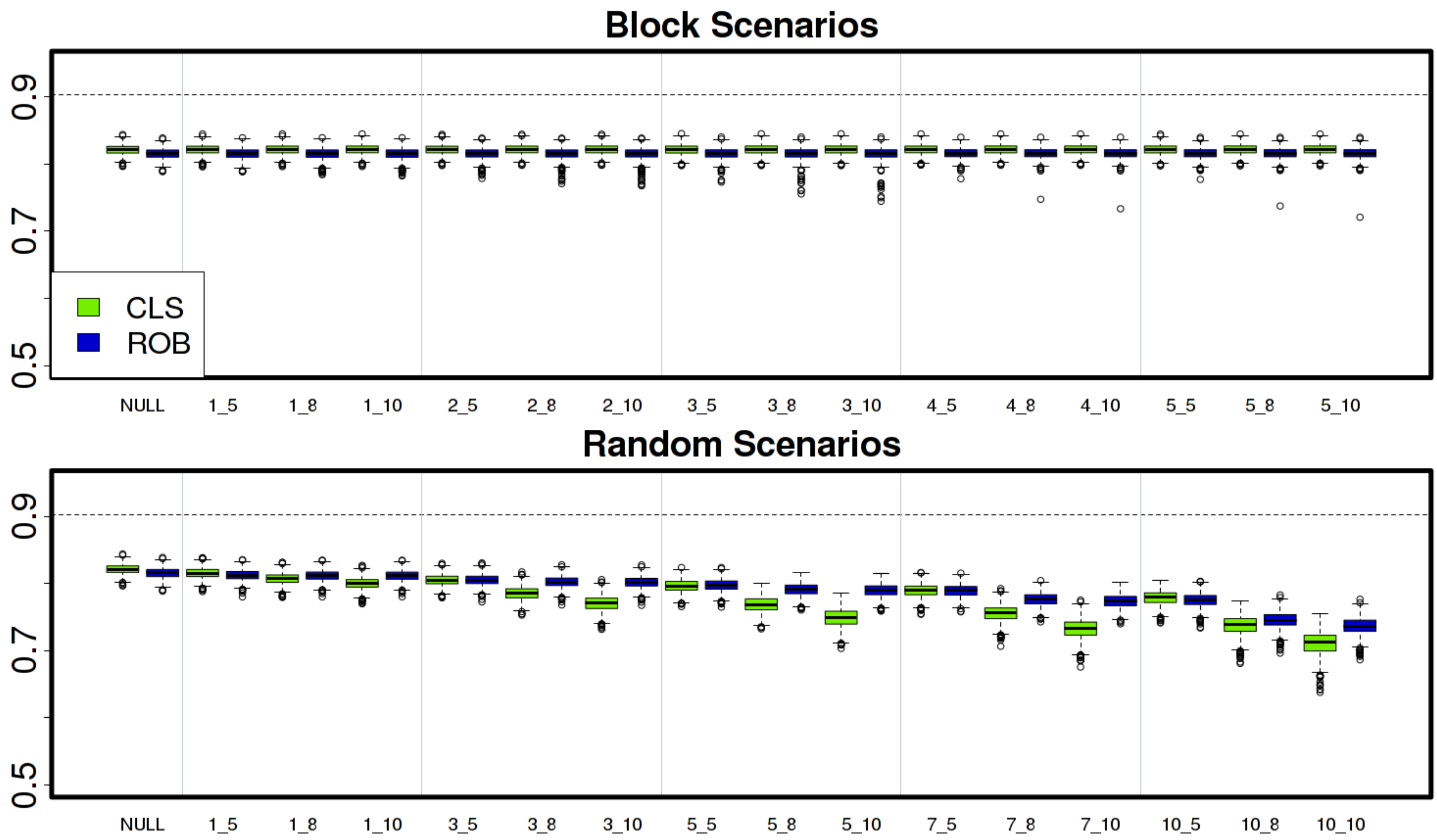


Figure 20S. Boxplots of estimated predictive accuracies (PA) computed by method **M7** for the **block** and **random** contamination scenarios (`3rd stage`)

### Appendix C

#### Robust Smith's weights and doubly robust Smith's weights

The robust estimation procedure weights each observation according to its relative contribution to the model fit while fitting the model at the first stage. In this way it attempts to explain most of the variation in the data. Here, we use simulation to investigate whether using doubly robust Smith's weights accounts for outliers better than using robust Smith's weights. The simulation considers 698 genotypes each with two replicates for a total of 1396 observations. The simulation encompasses three scenarios, namely the *null* (0% contamination), *block* 3_8 (3.5% contamination) and *random* 7_8 (3.9% contamination) scenarios, and considers 250 simulation runs for each scenario. Only 250 simulation runs are considered for each scenario because the simulations are computationally very expensive. The *block* 3_8 and *random* 7_8 simulation scenarios were chosen because (i) they have approximately the same percentage of overall contamination and (ii) the 8-sd shift-outliers are considered sufficiently extreme in size. We expect the weights computed for each of the two replicate observations to differ substantially if only one of the pair is an outlier.

The Smith's weights are the diagonal elements of the inverse of the estimated variance-covariance matrix from the first-stage model. Because the robust fit approximates most rather than all the data, the variances of the outlying observations will be greater than those for the regular observations. Therefore, the key idea of doubly robust Smith's weights is to account for this effect by re-weighting the Smith's weights from the first stage, by using either the minimum (*min*) or mean (*mean*) of the robust Smith's weights assigned to each pair of replicate observations in the first stage. We denote the standard robust Smith's weights with SW, robust Smith's weights re-weighted by the minimum ($SW_{min}$) or the mean ($SW_{mean}$) of the weights assigned to the two replicate observations at the first stage.

Figure C1 shows the differences in the distributions of the three different Smith's weights (SW, $SW_{min}$ and $SW_{mean}$) computed across the 250 simulation runs for the *null*, *block* 3_8 and *random* 7_8 scenarios. Notably, the weights for the *random* scenario are clearly smaller than those for either the null or block scenarios.

The mean squared deviations of $SW_{min}$ from the $SW_{mean}$ across the three scenarios and the 250 simulation runs is approximately $0.4 \times 10^{-11}$ for the *null* scenario, $0.2 \times 10^{-10}$ for the *block* 3_8 scenario and $0.7 \times 10^{-10}$ for the *random* 7_8 scenario. Figure C2 (first row) further corroborates the observation that the $SW_{min}$ and $SW_{mean}$ produce virtually the same results. However, there are noticeable differences between either of the re-weighted robust Smith's weights and the robust Smith's weights across all the three scenarios (Figure 2C, second-row). Overall, the robust Smith's weights are generally higher than the re-weighted ones. Furthermore, despite the MSDs between the robust Smith's weights and the re-weighted Smiths weights being numerically small, they are $10^5$ to $5 \times 10^5$ times greater than the MSDs of the Smith's weights re-weighted with *min* from those for the Smith's weights re-weighted with *mean*.

Figure C3, shows that for the 250 simulation runs for the *null*, *block* 3_8 and *random* 7_8 scenarios, there are no major differences between the estimated genetic variances obtained at the second-stage by using either the *min* or *mean* of the weights assigned to the two replicate observations to re-weight the robust Smith's weights from the first-stage model fit. In addition, the re-weighting scheme is most effective for the *random* contamination scenario for which using the robust Smith's weights alone is not sufficient to produce unbiased estimates of the true genetic variance.

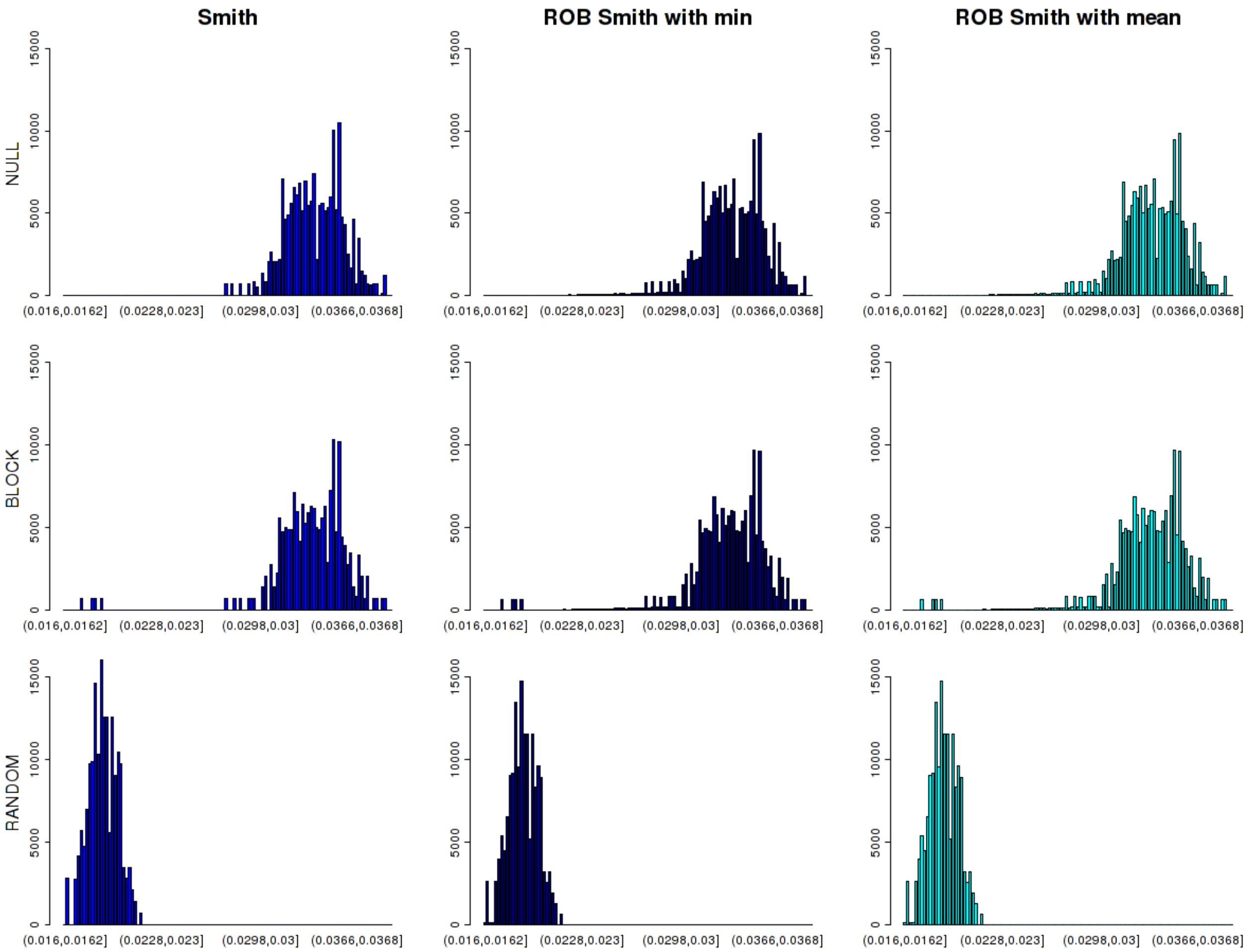


Figure C1: Frequency histograms of the Smith's weights computed from the first-stage robust model fit with and without re-weighting using the robust weights, for the *null*, *block* 3_8 and *random* 7_8 scenarios. The same number of bins and binwidths were used to facilitate comparison.

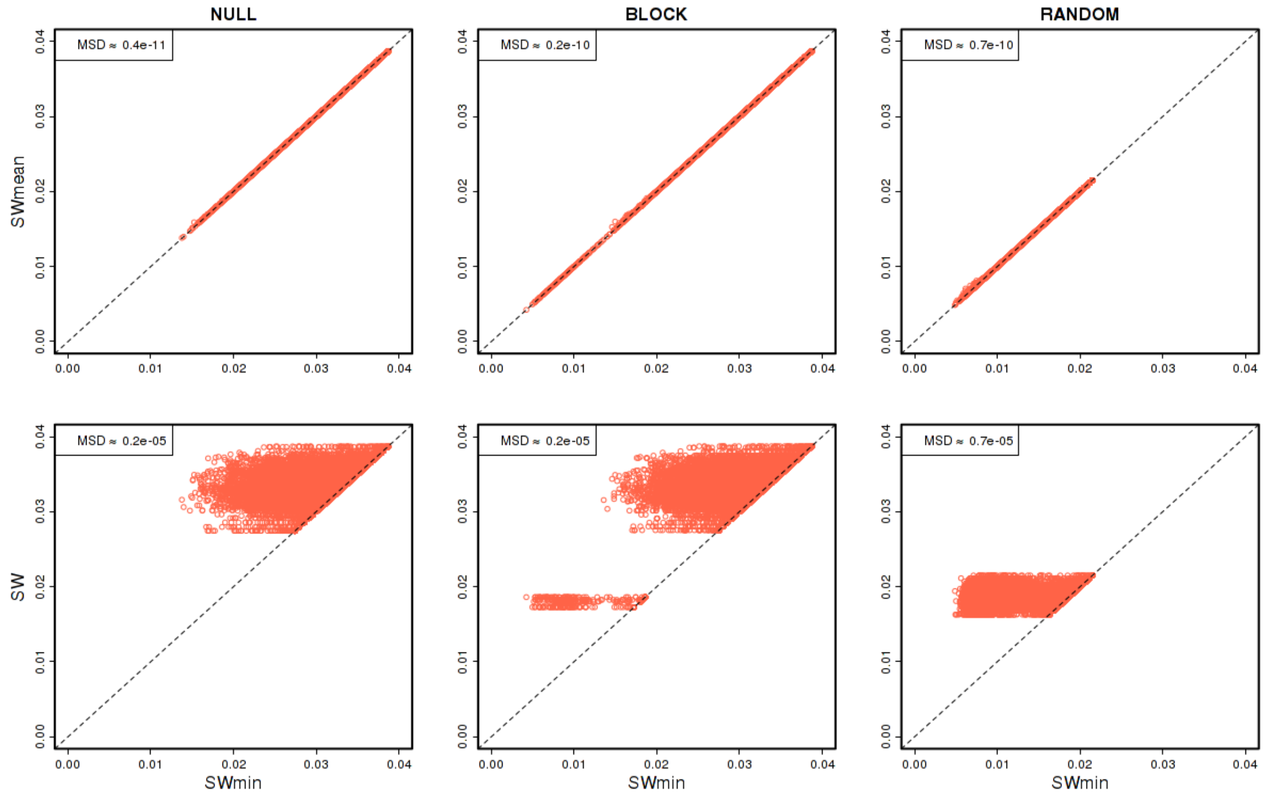


Figure C2: Plots of the Smith's weights computed from the first-stage robust fit with and without re-weighting using the robust weights, for the *null*, *block* 3_8 and *random* 7_8 scenarios. Re-weighted Smith's weights with *min* ($SW_{min}$) are plotted against the re-weighted Smith's weights with *mean* ($SW_{mean}$; first row) and re-weighted Smith's weights with *min* ($SW_{min}$) are plotted against the Smith's weights (SW).

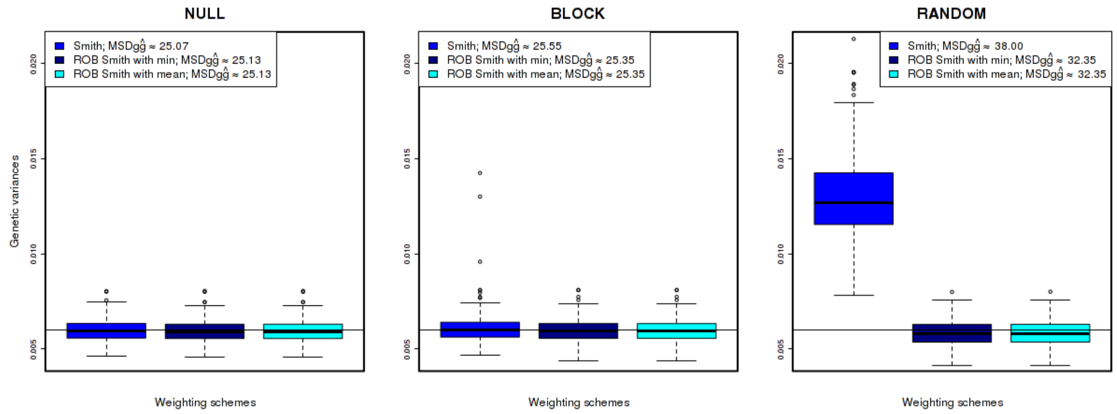


Figure C3: Boxplots of the 250 estimated robust genetic variances using the ordinary Smith's weights (as computed from the robust first-stage model; robust Smith's weights) and the weighted Smith's weights (computed by reweighting the Smith's weights with the robust weights obtained from the robust fit using both the minimum and the mean of the weights across the two replicates; doubly robust Smith's weights) for the *null*, *block* 3_8 and *random* 7_8 scenarios; Mean squared deviations (MSDs) from the estimated breeding values (EBVs) to the true breeding values (TBVs) are also shown.

### Appendix D

#### R code for the classical & robust two-stage approaches

```
## loading the data
  load("toydata.Rdata")

## loading the libraries
  library(lme4)         # for classical LMM
  library(robustlmm)    # for robust LMM
  library(asreml)       # for the classical LMM with kinship matrix
  library(psych)        # to being able of using the tr() function
  # if tr() does not work load library(matrixcalc) and use matrix.trace() function

## CLASSICAL first-stage fit below is commented;
## uncomment when you wish to run it, in which case you should comment the robust fit

# ## fitting the first-stage classical model
#   fit<-  lmer(yield ~ -1 + gen + (1|rep)+(1|rep:block), toydata)
#     # getting the lsmeans and var-covar structure
#       R   <- summary(fit)$vcov
#       mu  <- summary(fit)$coefficients[,1]
#     # computing the Smith's and Standard weights
#       w<-(1/diag(R))           #Tandard weights
#       wsmith<-diag(solve(R))   #Smith's weights
#       rm(R)
#     # keeping the estimated random effect variances
#       STDs<-matrix(0,3,1)
#       STDs[1,1]<- attr(VarCorr(fit)$'rep:block', "stddev")
#       STDs[2,1]<- attr(VarCorr(fit)$'rep', "stddev")
#       STDs[3,1]<- attr(VarCorr(fit),"sc")
#       colnames(STDs)<-"std"
#       rownames(STDs)<-c("REP:BLOCK","REP","Residual")
#       stage1.vars<-STDs^2
#       rm(STDs)

## fitting the first-stage robust model
  fit<-  rlmer(yield ~ -1 + gen + (1|rep)+(1|rep:block), toydata,
             rho.sigma.e = psi2propII(smoothPsi, k = 2.28))
    # getting the lsmeans and var-covar structure
      R  <- summary(fit)$vcov
      mu <- summary(fit)$coefficients[,1]
    # getting the robust weights
    # do not confuse these with the Smith's and Standard weights
      rob.weights<-getME(fit,name="w_e")
    # the robust weights need not be the same for genos in rep1 and rep2
    # but we want only 1 robust weight per-genotype
    # thus we will choose the min between the 2 robust weights from the 2 replicates
    # The next computations need to be adapted for each dataset because the order of
    # the weights matches the one for one of the datasets as does also the order of the residuals
      n<-length(ls.means)
      aux<-vector()
      for(k in seq(1,(n*2-1),by=2)){aux<-c(aux,min(rob.weights[k],rob.weights[k+1]))}
      rob.weights<-aux
```

```
      rm(aux,k,n)
    # computing the Smith's and Standard weights, which incorporate the robust weights
      w     <-(1/diag(R))*rob.weights       #Standard weights
      wsmith<-diag(solve(R))*rob.weights    #Smith's weights
      rm(rob.weights,R)
    # keeping also the estimated random effects variances
      STDs<-matrix(0,3,1)
      STDs[1,1]<- attr(VarCorr(fit)$'rep:block', "stddev")
      STDs[2,1]<- attr(VarCorr(fit)$'rep', "stddev")
      STDs[3,1]<- attr(VarCorr(fit),"sc")
      colnames(STDs)<-"std"
      rownames(STDs)<-c("REP:BLOCK","REP","Residual")
      stage1.vars<-STDs^2
      rm(STDs)

## fitting the second-stage model -- classical approach used
  # # try out G=I to see how H2.M5 and H2.Oakey match
  #   toyG<-diag(dim(toyG)[1])
  #   colnames(toyG)<-names(mu)
  #   rownames(toyG)<-names(mu)

  # preparing the data
    plantid<-names(mu)
    colnames(toyG)<-names(mu)
    rownames(toyG)<-names(mu)
    inv.toyG<-solve(toyG)
    ourdata<-data.frame(plantid = plantid,
                        mu = mu,
                        wsmith = wsmith,
                        w = w)
  # fitting the model
  # one can replace the Smith's weights (wsmith) with the Standard weights (w) below
    fit.cls <-asreml( data    = ourdata,
                      fixed   = mu ~ 1,
                      random  =~ giv(plantid) ,
                      rcov    =~ units, na.method.Y = "include",
                      weights = wsmith,
                      family  = asreml.gaussian(dispersion=1.0),
                      control = asreml.control(workspace=16e7, ginverse=list(plantid=inv.toyG),
                                        maxiter=1000)
                    )
    # computing the eBLUPs, estimated genetic variance and C22 matrix
      gBLUP <-fit.cls$coefficients$random
      g.var <-summary(fit.cls)$varcomp['giv(plantid).giv','component']
      C22   <- predict(fit.cls, classify="giv(plantid)", only="giv(plantid)", vcov=T)$pred$vcov
    # removing stuff from memory
      rm(fit,fit.cls,ourdata,toydata,plantid)
      rm(inv.toyG)

## third-stage -- Estimating heritability and predictive accuracy
  # Preparing the matrices and auxiliary variables using the notation in the paper
    G      <-toyG
    n      <-dim(G)[1]
```

```
  G.tilde<-G*g.var
  R.tilde<-solve(diag(n)*(wsmith))
  rm(toyG)

# computing heritability and predictive accuracy via METHOD 5
  V    <-G.tilde+R.tilde
  P    <-(1/(n-1))*(diag(n)-matrix(1,n,n)/n)
  one  <-as.matrix(rep(1,n))
  Q    <-diag(n)-one %*% solve(t(one)%*%solve(V)%*%one) %*% t(one) %*% solve(V)
  C    <-G.tilde%*%solve(V)%*%Q

  PA.est.m5<-tr(P%*%C%*%G.tilde)/sqrt(tr(P%*%G.tilde)*tr(t(C)%*%P%*%C%*%V))
  H2.est.m5<-PA.est.m5^2
  rm(V,P,Q,C,one)

# computing reliability and predictive accuracy via METHOD 7
  v1<-G.tilde
  v2<-G.tilde-C22
  rho2<-vector()
  for(j in 1:n){rho2[j]<-(v2[j,j])^2/(v1[j,j]*v2[j,j])}
  rm(j,v1,v2)

  RL.est.m7       <-mean(rho2)
  PA.est.m7       <-mean(sapply(rho2,sqrt))
  rm(rho2)

# computing heritability via OAKEY's METHOD
  D     <-diag(n)-solve(G.tilde)%*%C22
  eival <-eigen(D)$values
  s     <-length(eival[eival<0.0001])

  H2.OAKEY<-tr(D)/(n-s)
  rm(D,eival,s,G.tilde,R.tilde)

  rm(n,w,wsmith)
  rm(G,C22)

  cbind(H2.est.m5,H2.OAKEY,PA.est.m5,PA.est.m7)
```
